# Spatially-resolved correlative microscopy and microbial identification reveals dynamic depth- and mineral-dependent anabolic activity in salt marsh sediment

**DOI:** 10.1101/2020.08.03.234146

**Authors:** Jeffrey Marlow, Rachel Spietz, Keun-Young Kim, Mark Ellisman, Peter Girguis, Roland Hatzenpichler

## Abstract

Coastal salt marshes are key sites of biogeochemical cycling and ideal systems in which to investigate the community structure of complex microbial communities. Here, we clarify structural-functional relationships among microorganisms and their mineralogical environment, revealing previously undescribed metabolic activity patterns and precise spatial arrangements within salt marsh sediment. Following 3.7-day *in situ* incubations with a non-canonical amino acid that was incorporated into new biomass, samples were embedded and analyzed by correlative fluorescence and electron microscopy to map the microscale arrangements of anabolically active and inactive organisms alongside mineral grains. Parallel sediment samples were examined by fluorescence-activated cell sorting and 16S rRNA gene sequencing to link anabolic activity to taxonomic identity. Both approaches demonstrated a rapid decline in the proportion of anabolically active cells with depth into salt marsh sediment, from ∼60% in the top cm to 10-25% between 2-7 cm. From the top to the bottom, the most prominent active community members shifted from sulfur cycling phototrophic consortia, to sulfate-reducing bacteria likely oxidizing organic compounds, to fermentative lineages. Correlative microscopy revealed more abundant (and more anabolically active) organisms around non-quartz minerals including rutile, orthoclase, and plagioclase. Microbe-mineral relationships appear to be dynamic and context-dependent arbiters of biogeochemical cycling.

**Statement of Significance:** Microscale spatial relationships dictate critical aspects of a microbiome’s inner workings and emergent properties, such as evolutionary pathways, niche development, and community structure and function. However, many commonly used methods in microbial ecology neglect this parameter – obscuring important microbe-microbe and microbe-mineral interactions – and instead employ bulk-scale methodologies that are incapable of resolving these intricate relationships.

This benchmark study presents a compelling new approach for exploring the anabolic activity of a complex microbial community by mapping the precise spatial configuration of anabolically active organisms within mineralogically heterogeneous sediment through *in situ* incubation, resin embedding, and correlative fluorescence and electron microscopy. In parallel, active organisms were identified through fluorescence-activated cell sorting and 16S rRNA gene sequencing, enabling a powerful interpretive framework connecting location, identity, activity, and putative biogeochemical roles of microbial community members.

We deploy this novel approach in salt marsh sediment, revealing quantitative insights into the fundamental principles that govern the structure and function of sediment-hosted microbial communities. In particular, at different sediment horizons, we observed striking changes in the proportion of anabolically active cells, the identities of the most prominent active community members, and the nature of microbe-mineral affiliations. Improved approaches for understanding microscale ecosystems in a new light, such as those presented here, reveal environmental parameters that promote or constrain metabolic activity and clarify the impact that microbial communities have on our world.

## Introduction

Salt marshes are vibrant microbial habitats that play important roles in the biogeochemical cycling of intertidal ecosystems (Tobias and Neubauer, 2019). The confluence of high organic input and seawater-derived sulfate fuel a wide range of carbon, nitrogen, phosphorous, and sulfur transformations over compressed spatial scales, leading to abundant, redox-specific niches and microbial communities with high phylogenetic diversity (Lozupone and Knight, 2007; Bowen *et al*., 2012). Because of this, salt marshes represent ideal sites to explore the intricacies of microbial community structure from the microscale to the ecosystem scale.

Within complex microbial communities, spatial relationships are increasingly seen as central determinants of key ecological parameters. In salt marshes, metabolic activity within specific sediment horizons ultimately shapes emergent properties such as carbon sequestration or greenhouse gas emissions (Abdul-Aziz *et al*., 2018; LaRowe *et al*., 2020). More generally, microbe-microbe and microbe-mineral interactions establish evolutionary trajectories (Cordero *et al*., 2012; Andersen *et al*., 2015), niche development (Morton *et al*., 2017), and community structure, function, and stability (Boetius *et al*., 2000; Wright *et al*., 2012; Coyte *et al*., 2015). Further, inter-organism arrangements govern chemical communication (West *et al*., 2007), metabolite exchange (Romine *et al*., 2017), and competition for resources (Mitri *et al*., 2016). Nonetheless, these critical spatial relationships are neglected by the most commonly used methods in microbial ecology, such as bulk meta-omics and geochemical approaches. As a result, important metabolic activities may be obscured, including inter-species nutrient cycling (Wilbanks *et al*., 2014; Cordero and Datta, 2016) and electron transfer to (Lovley and Phillips, 1988; Myers and Nealson, 1988) or from (Shelobolina *et al*., 2012) specific minerals.

Recent efforts have made progress in analyzing microbial communities at the microscale. nanoscale secondary ion mass spectrometry (nanoSIMS) coupled with stable isotope probing (SIP) and fluorescence *in situ* hybridization (FISH) can resolve anabolic patterns and taxonomically identify individual cells. However, this method typically separates microbial assemblages from their broader environmental context (McGlynn *et al*., 2015; Musat *et al*., 2016; Gyngard and Steinhauser, 2019). By combining energy dispersive x-ray spectroscopy (EDS) with x-ray computed tomography images, Hapca *et al*. extended chemical analyses into a third dimension with resin-embedded soil, but no cellular information was attained (Hapca *et al*., 2015). Correlative imaging with nanoSIMS and electron and fluorescence microscopy enabled Schlüter *et al*. to pinpoint the position of a subset of the microbial community in relation to leaf fragments, but metabolic activity and microbial identities were not considered (Schlüter *et al*., 2018). A promising addition to this emerging field is SIP combined with non-destructive Confocal Raman microspectroscopy, which was recently used to measure the *in situ* activity and substrate uptake of microbes in transparent soil microcosms (Sharma *et al*., In Press).

The work presented here advances this line of microbial ecology research. The methods herein not only preserve spatial arrangements and link cell positions to mineralogy through correlative microscopy, but also establish the presence, location, and mineralogical associations of anabolically active cells. Anabolic activity was assessed with bioorthogonal non-canonical amino acid tagging (BONCAT), a next-generation physiology approach (Hatzenpichler *et al*., 2020) that uses substrate analog probing to visualize protein synthesis in active cells. A non-canonical amino acid, such as *L*-homopropargylglycine (HPG) or *L*-azidohomoalanine (AHA), is incorporated into growing peptides by native methionyl-tRNA synthetases. Subsequent azide-alkyne click chemistry allows fluorescent detection of newly synthesized proteins (Sletten and Bertozzi, 2009). BONCAT was initially developed in neuron (Dieterich *et al*., 2006), eukaryote (Hinz *et al*., 2011), and cultured bacteria (Hatzenpichler *et al*., 2014) systems; more recently, it was optimized for environmental microbial communities and shown to have no measurable effect on community composition or metabolic activity (Hatzenpichler *et al*., 2014, 2016). The approach has been proven effective in a diverse range of bacterial and archaeal cultures (Hatzenpichler *et al*., 2014; Hatzenpichler and Orphan, 2015); ocean water (Samo *et al*., 2014; Leizeaga *et al*., 2017; Sebastián *et al*., 2019), marine sediment (Hatzenpichler *et al*., 2016), hot spring (Reichart *et al*., 2020), and soil microbiomes (Couradeau *et al*., 2019); as well as marine viruses and bacteriophages (Pasulka *et al*., 2018). BONCAT appears to be a taxonomically agnostic measure of anabolic activity that correlates well with other metrics of activity (Bagert *et al*., 2014; Hatzenpichler *et al*., 2014, 2020) with only small effects on metabolism (Steward *et al*., 2020) and protein chemistry (Bagert *et al*., 2014; Lehner *et al*., 2017).

In this study, we mapped the anabolic activity of individual microorganisms in sediments from Little Sippewissett Salt Marsh (LSSM) in Falmouth, MA. In the LSSM, terrestrial freshwater runoff, seawater, high organic input, and abundant light and chemical energy leads to dramatic redox stratifications within the top few centimeters of sediment and a wide range of metabolic niches (Armitage *et al*., 2012; Wilbanks *et al*., 2014, 2017; Larsen *et al*., 2015). Using purpose-built equipment, a series of sediment cores were incubated with HPG *in situ* for 3.7 days. One set of cores was used for correlative microscopy; samples were embedded in resin to maintain precise spatial arrangements, sectioned, stained, and analyzed using fluorescence and electron microscopy to map active and inactive biomass as well as identifiable mineral grains. A parallel set of cores was processed for horizon-specific fluorescence activated cell sorting (FACS) and 16S rRNA gene amplicon sequencing. With this novel approach, we mapped active and inactive organisms in their native microscale configuration and identified the active and inactive microbial communities in adjacent sediment horizons.

Our results indicate that the proportion of anabolically active organisms decreased dramatically below the photic zone, and that mineralogy likely has an impact on the relative abundance and anabolic activity of mineral grain-associated organisms. High-throughput 16S rRNA gene sequencing of active and inactive microbial communities in adjacent sediment cores revealed a continuous progression of community structure with depth, oriented around shifting metabolisms of photosynthesis, sulfur cycling, and fermentation. Notably, with correlative fluorescence and electron microscopy, we observed differential cell association with distinct mineral types and a greater proportion of organisms inside mineral grains in lower (6-7 cm) sediment horizons compared with shallower zones. While the full potential of microbiome mapping remains to be realized, this benchmarking study unveils a new experimental approach to a) evaluate how metabolic activity relates to microscale environmental factors, and b) develop testable hypotheses regarding metabolic interactions among members of complex microbial communities.

## Results & Discussion

This study reveals how microbial presence and anabolic activity relate to mineralogical distributions at the microscale with a new level of realism in salt marsh sediment. Correlative microscopy analyses at three distinct horizons revealed changes in organism abundance from 1.95×10^9^ cm^−3^ at 7.6 mm depth to 2.86×10^9^ cm^−3^ at 12 mm depth and 6.85×10^8^ cm^−3^ at 60.7 mm depth. Moving downward along these three horizons, the proportion of anabolically active organisms decreased from 51.3% (7.6 mm) to 22.3% (12 mm) to 12.1% (60.7 mm), a trend that correlated well with BONCAT-FACS data (R^2^=0.99; Table 1).

**Table 1:**
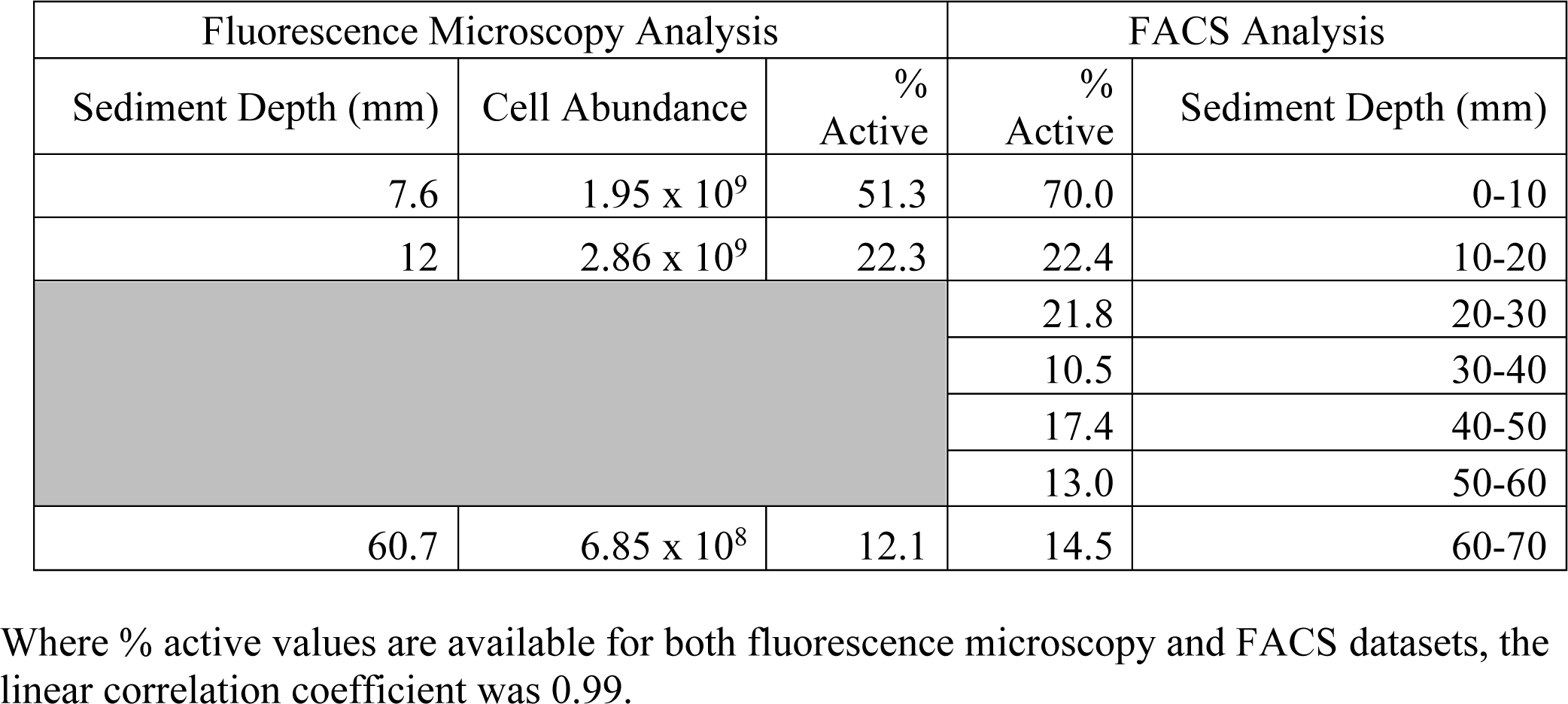
Cell abundance and percentage of anabolically active cells as determined through fluorescence microscopy and BONCAT-FACS analyses.

| Fluorescence Microscopy Analysis |  |  | FACS Analysis |  |
| --- | --- | --- | --- | --- |
| Sediment Depth (mm) | Cell Abundance | % Active | % Active | Sediment Depth (mm) |
| 7.6 | $1.95 \times 10^9$ | 51.3 | 70.0 | 0-10 |
| 12 | $2.86 \times 10^9$ | 22.3 | 22.4 | 10-20 |
|  |  |  | 21.8 | 20-30 |
|  |  |  | 10.5 | 30-40 |
|  |  |  | 17.4 | 40-50 |
|  |  |  | 13.0 | 50-60 |
| 60.7 | $6.85 \times 10^8$ | 12.1 | 14.5 | 60-70 |

Where % active values are available for both fluorescence microscopy and FACS datasets, the linear correlation coefficient was 0.99.

At each microscopy horizon, the mineralogical identities of individual grains were assessed in order to determine whether different mineral types corresponded with notable differences in organism abundance, configuration, or anabolic activity. The majority of all detected grains were quartz (SiO_2_), while albite (NaAlSi_3_O_8_), orthoclase (KAlSi_3_O_8_), rutile (TiO_2_), plagioclase (a solid solution range from NaAlSi_3_O_8_ to CaAl_2_Si_2_O_8_), and Ca/K/Mg/Fe silicate grains of indeterminant mineralogy were also detected. Like the vocabulary used to describe microbe-microbe interactions, microbe-mineral interactions can be harmful, neutral, or beneficial for the organism. Microbial sorption to quartz grains has been demonstrated, but repellant electrical charges make the interaction less favorable than those with other mineral types (Mills *et al*., 1994; Gong *et al*., 2018). The best-studied beneficial interactions are the microbial reduction of iron or manganese oxides (Thamdrup, 2000), which enables bacteria to off-load reducing power, altering mineral structure and chemistry in the process (Kawano and Tomita, 2002; Welch and Banfield, 2002). A number of factors influence the nature of these interactions, including accessible surface area, mineral lattice structure, co-occurrence of organic matter, and other environmental conditions such as temperature and pH (Dong *et al*., 2009). Beyond iron and manganese, microbes have been shown to associate with other cations, acquiring potassium from silicates (Valsami-Jones *et al*., 1998), releasing organic ligands that adhere to aluminum (Rogers and Bennett, 2004), and using reducing power from photo-catalytically activated titanium oxide (Lu *et al*., 2012).

### Top horizon BONCAT-FACS & Correlative Microscopy

Many previous studies have elucidated key aspects of the LSSM microbiological system and its role in biogeochemical cycling (Seitz *et al*., 1993; Shapiro *et al*., 2011; Armitage *et al*., 2012; Bowen *et al*., 2013; Peng *et al*., 2013; Wilbanks *et al*., 2014, 2017; Larsen *et al*., 2015; Mackey *et al*., 2017; Angell *et al*., 2018); we leverage this heritage to infer physiological traits based on the 16S rRNA gene data we collected. (Please see Supporting Information Dataset 1 for sequence data and relative abundances of assigned lineages for bulk, active, and inactive samples across all horizons.)

The top ten millimeters of LSSM sediment exhibit dramatic redox gradients as oxygen concentrations fall below detection by 5 mm, sulfide rises from 0 to between 0.5-1.5 mM, and pH fluctuates between ∼7.0-7.3 at night and ∼6.0-7.0 during the day (Armitage *et al*., 2012; Larsen *et al*., 2015; Salman *et al*., 2015). The microbial community was dominated by the phyla *Proteobacteria* (48% relative abundance) and *Bacteroidetes* (30%), whose metabolically diverse members are indicative of a range of redox conditions and substantial heterotrophic cycling in the upper sediment layer (Spain *et al*., 2009; Gómez-Pereira *et al*., 2012). *Thiohalocapsa* was the most abundant genus-level lineage recovered, accounting for 14.4% of all sequences; *Desulfobulbaceae* was the next most abundant genus, with two unidentified lineages representing 3.8% and 2.7% of all sequences. The prevalence of these purple sulfur bacteria and sulfate-reducing bacteria is reflective of the abundant “pink berries” found at the sediment surface (Fig. S1) (Seitz *et al*., 1993; Wilbanks *et al*., 2014). Among organisms putatively involved with sulfur-cycling consortia, we observed a more diverse distribution of sulfate-reducing bacteria lineages (65 genus-level *Desulfobacterales* ASVs) and a more streamlined set of purple sulfur bacteria with a single dominant representative (19 genus-level *Chromatiales* ASVs, with *Thiohalocapsa* accounting for 83% of the recovered sequences).

During the 3.7-day incubation period, the majority of organisms detected in this sediment zone demonstrated anabolic activity (Table 1). Sequencing of active and inactive communities in the 0-10 mm range revealed eight lineages representing >1% of the overall relative abundance that were significantly more abundant in the anabolically active subset (Dataset 1). Of these, six were putative members of the pink berry consortia (*Chromatiales* or *Desulfobacterales* orders), one was a photoheterotroph that may encode multiple light-harvesting complexes (*Halieaceae*, (Spring *et al*., 2015)), and one was a representative of the metabolically diverse *Rhodobacteraceae* family (Pujalte *et al*., 2014; Pohlner *et al*., 2019). Many of the other abundant inactive lineages – including three putative sulfate reducers and three putative purple sulfur bacteria – were among the most abundant ASVs in both the active and inactive fractions (Dataset 1). This overlap may indicate stochastic activity of particular consortia or a metabolic dependence upon physicochemical traits on a sub-cm scale, such as pore connectivity or identity of neighboring organisms. Alternatively, our conservative gating approach may have captured some active cells with low fluorescence in the inactive gate (Fig S2). Among the inactive microorganisms, two *Rhizobiaceae* lineages constituted a combined 11.8% of the sequenced fraction. This family of *Alphaproteobacteria* is typically associated with actively growing plants (Spaink *et al*., 2012); their anabolic quiescence could be attributable to displacement from spartina grass roots surrounding the sample site.

The uppermost section examined by correlative microscopy was located within the top sequenced horizon, at a depth of 7.6 mm (Fig. 1). In the analysis area, 15 of the 20 mineral grains were quartz (SiO_2_), while albite (NaAlSi_3_O_8_), orthoclase (KAlSi_3_O_8_), rutile (TiO_2_), plagioclase (a solid solution range from NaAlSi_3_O_8_ to CaAl_2_Si_2_O_8_), and Ca/K/Mg/Fe silicate grains of indeterminant mineralogy were also observed. 73.4% of cells were located within 70 µm or found inside of quartz grains. When cell biomass abundances were normalized by proxies for mineral surface area and volume, non-quartz grains exhibited a higher organism density (Table 2).

**Fig. 1:**
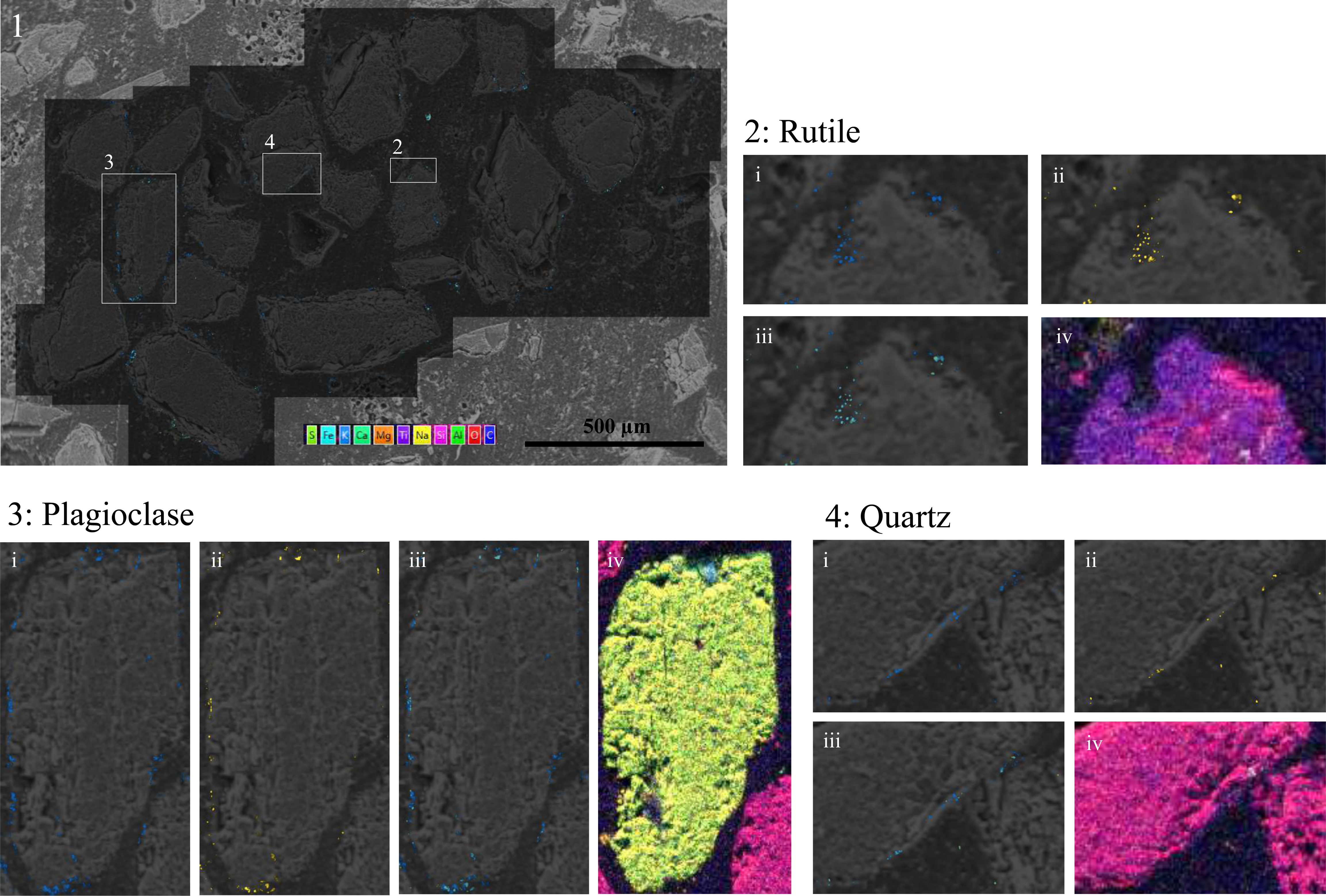
Correlative fluorescence and electron microscopy from the uppermost section (7.6 mm sediment depth). 1) Overlain on the base SEM image are two fluorescence channels showing SYBR-active features in blue, and BONCAT-active features in yellow. The dark zonation indicates the fluorescence microscopy footprint. 2), 3), and 4) show three mineralogically distinct sites in detail. i) SYBR green, ii) BONCAT, and iii) merged channels, as well as iv) EDS elemental abundance maps (in which dark blue background represents the resin).

**Table 2:**
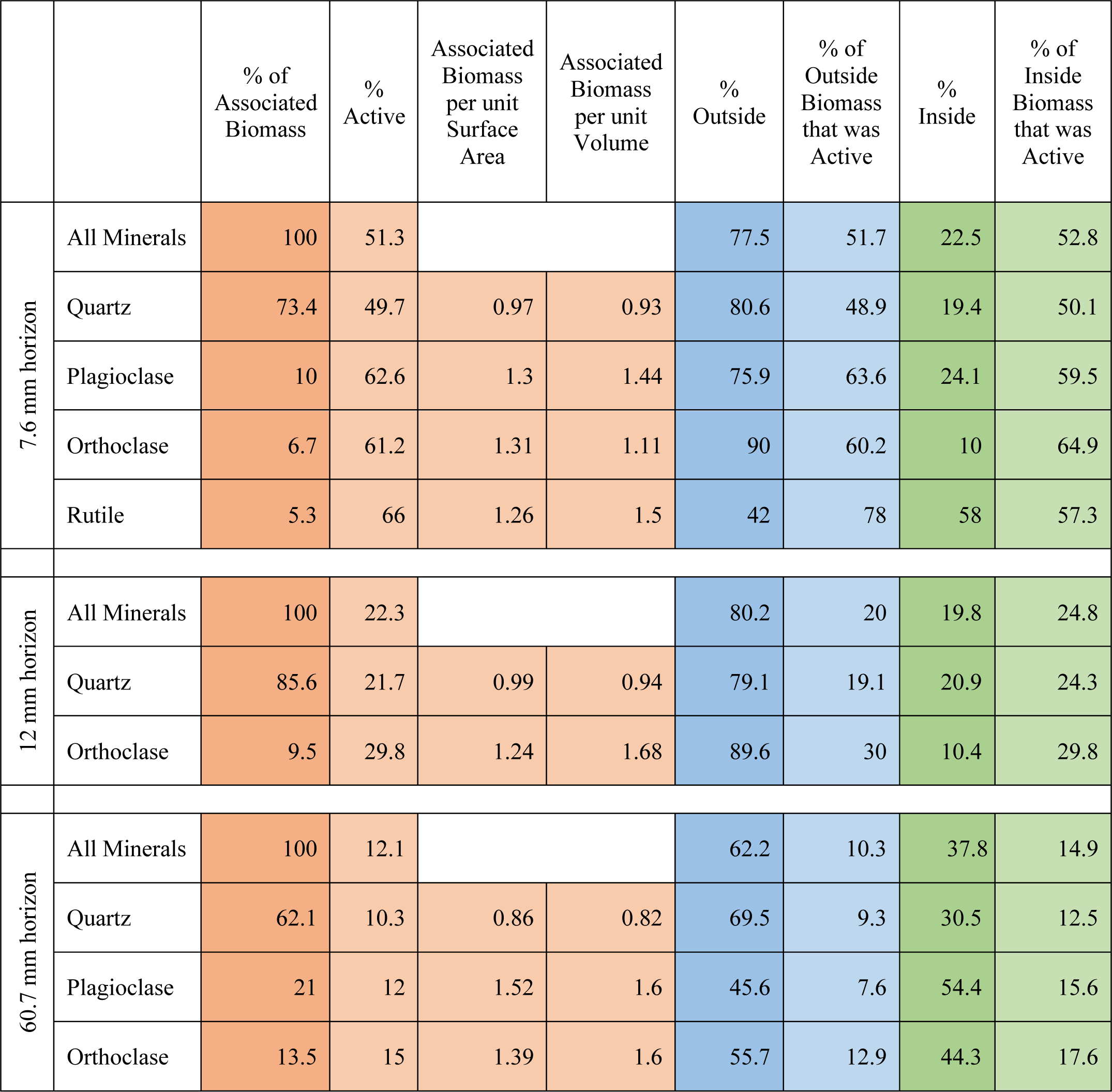
Proportions of cells, and the anabolically active subsets, associated with mineral exteriors and interiors at the three horizons examined by correlative microscopy. For the biomass per surface area and volume, the relative proportion of biomass associated with a given mineral type was divided by the relative proportion of surface area or volume accounted for by that mineral type. Values less than 1 indicate fewer associated cells than would be expected given an even distribution of biomass across mineral perimeters or surfaces. Only mineral types that accounted for at least 5% of the observed biomass in a given horizon are included in this analysis.

Overall, 77.5% of observed cells were outside their associated mineral grains while 22.5% were found inside, frequently along fractures or pores visible by SEM. The most biomass-rich zone was the “surface-associated” 0-5 µm bin, where 27.8% of cells were found. Beyond 10 µm, cell abundance decreased markedly, demonstrating a strong preference for grain interface zones and reflecting the removal of suspended biomass when the original porewater was replaced with filtered HPG solution prior to incubation. These spatial trends were broadly consistent across different mineral types (Fig. 2), though the greatest proportion (39%) of orthoclase-associated cells were not in the “surface-associated” 0-5 µm bin, but in the 5-10 µm zone. We also found that the degree of anabolic activity was higher around non-quartz minerals when compared with quartz-associated cells (Table 2). Although low abundances of these mineral types make generalizations difficult, it is possible that metal cations in the mineral structures facilitate a wider range of metabolic reactions than the more chemically inert quartz (Shi *et al*., 2016). The electrical semi-conductivity of titanium oxide can promote extracellular electron transfer (Zhou *et al*., 2018) and, via photo-catalysis, stimulate the growth of non-phototrophic microbes (Lu *et al*., 2012); these mechanisms may account for the elevated proportion (78%, compared with a mean of 51.7% for this horizon) of active cells associated with the exterior of the titanium oxide rutile grain.

**Fig 2:**
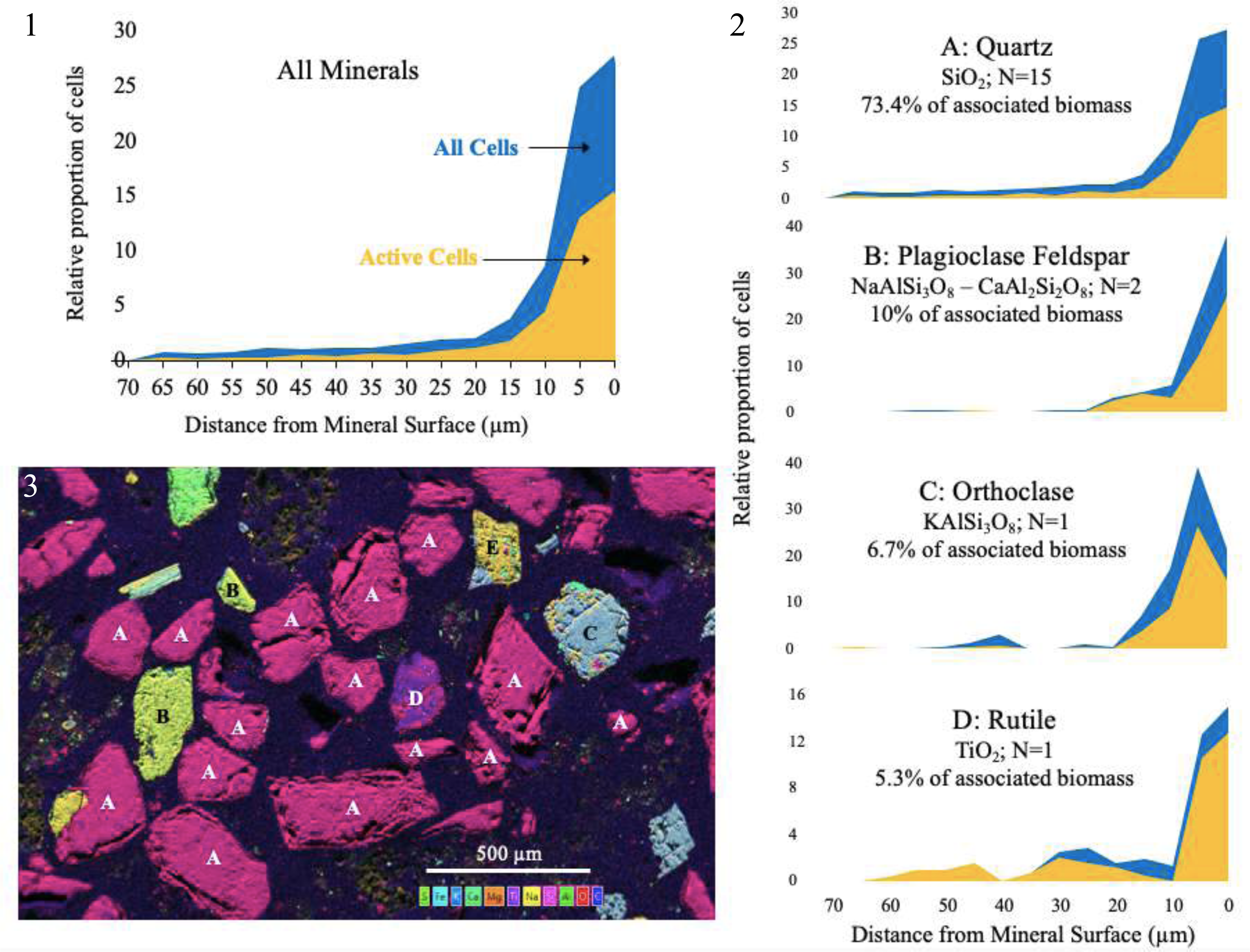
Histograms of the relative proportions of all organisms and the anabolically active subset (yellow overlay) at given distances from the mineral surface for the uppermost section (7.6 mm sediment depth). 1) Data for all grains. 2) Data separated by mineral type. Histogram bins are in 5 µm intervals, and only cells located outside mineral surfaces are shown. 3) Composite elemental maps derived from EDS analysis show the mineral grains that were analyzed, labeled by mineral type. A = quartz; B = Plagioclase; C = Orthoclase; D = Rutile; E = Albite; F = Ca,K,Mg,Fe silicate; G = Hornblende.

### Second horizon BONCAT-FACS & Correlative Microscopy

Between 10-20 mm of sediment depth, oxygen is absent and pH remains largely consistent within a ∼0.3 unit range, but sulfide concentrations exhibit substantial (up to ∼500 µM) fluctuations upward or downward, with no clear pattern based on diurnal cycle timing (Larsen *et al*., 2015; Salman *et al*., 2015). Under these more energetically constrained conditions, 16S rRNA gene reads were dominated by *Proteobacteria* (40%), *Cyanobacteria* (26%), and *Bacteroidetes* (13%) phyla suggestive of recycling of plant material as well as vibrant nitrogen and sulfur cycling processes. Among the ASVs representing more than 1% of relative abundance, one ASV most similar to mitochondria (2.8%) and eight cyanobacterial chloroplast-like sequences (summing to 16.7%) likely reflect burial and degradation of animal and photosynthetic biomass. Primers used in this study can amplify some eukaryotic sequences (Parada *et al*., 2016), but the short sequence length precludes accurate taxonomic assignment to eukaryotic chloroplast or mitochondria. Interestingly, these constituents were seemingly selected against during the cell extraction and sorting process, as their relative abundance in the active and inactive fractions were all well below 1%. Mitochondrial ASVs accounted for 0.12% and 0.01% of sequences in active and inactive fractions, respectively; chloroplast ASVs were 0% and 0.01%, respectively.

The contribution of burial and degradation is consistent with the lower proportion of active organisms observed by both microscopy (22.3%) and cell sorting data (22.4%). Sulfate-reducing deltaproteobacterial clades (particularly those within the *Desulfobacterales* family) as well as the environmentally wide-ranging betaproteobacterial family *Burkholderiaceae*, were more prevalent among active organisms. Sulfate-reducing bacteria represented five of the seven ASVs that were significantly enriched in the active fraction and accounted for at least 1% of the overall relative abundance, indicating a vibrant sulfur cycle likely fueled by organic carbon degradation. *Rhizobiales* and *Chromatiaceae* were more abundant in the inactive fraction, suggesting that potentially critical environmental factors like viable plant cells and sunlight, respectively, were not abundantly available. Nonetheless, one *Chromatiaceae* ASV (of the *Halochromatium* genus) was the second most-abundant lineage among active organisms, indicating that anoxygenic photosynthesis was still possible at this sediment depth (and/or bioturbation contributed to in-mixing from more photosynthetically active surface layers).

Within the 10-20 mm depth zone, a post-BONCAT embedded section from a depth of 12 mm was examined by correlative microscopy (Figs. 3-4). Twenty-two mineral grains were analyzed; as above, the vast majority of grains were quartz, and the microbes associated with non-quartz grains (in particular, orthoclase) had a higher proportion of anabolically active constituents (Table 2). Across all mineral types, exterior organisms were more spatially constrained to surfaces than the analyzed section from 7.6 mm: 40.5% were located within 5 µm of mineral grains compared to 27.8% in the top layer. Some of the highest concentrations of active cells were associated not with well-defined minerals, but rather with heterogeneous patches that include small particles of quartz, sodium, and iron (Fig. 3). In comparison with larger mineral grain interfaces, these particle assemblages offer greater chemically diversity and more potentially reactive surface area, factors that may facilitate interactions among microbes.

**Fig. 3:**
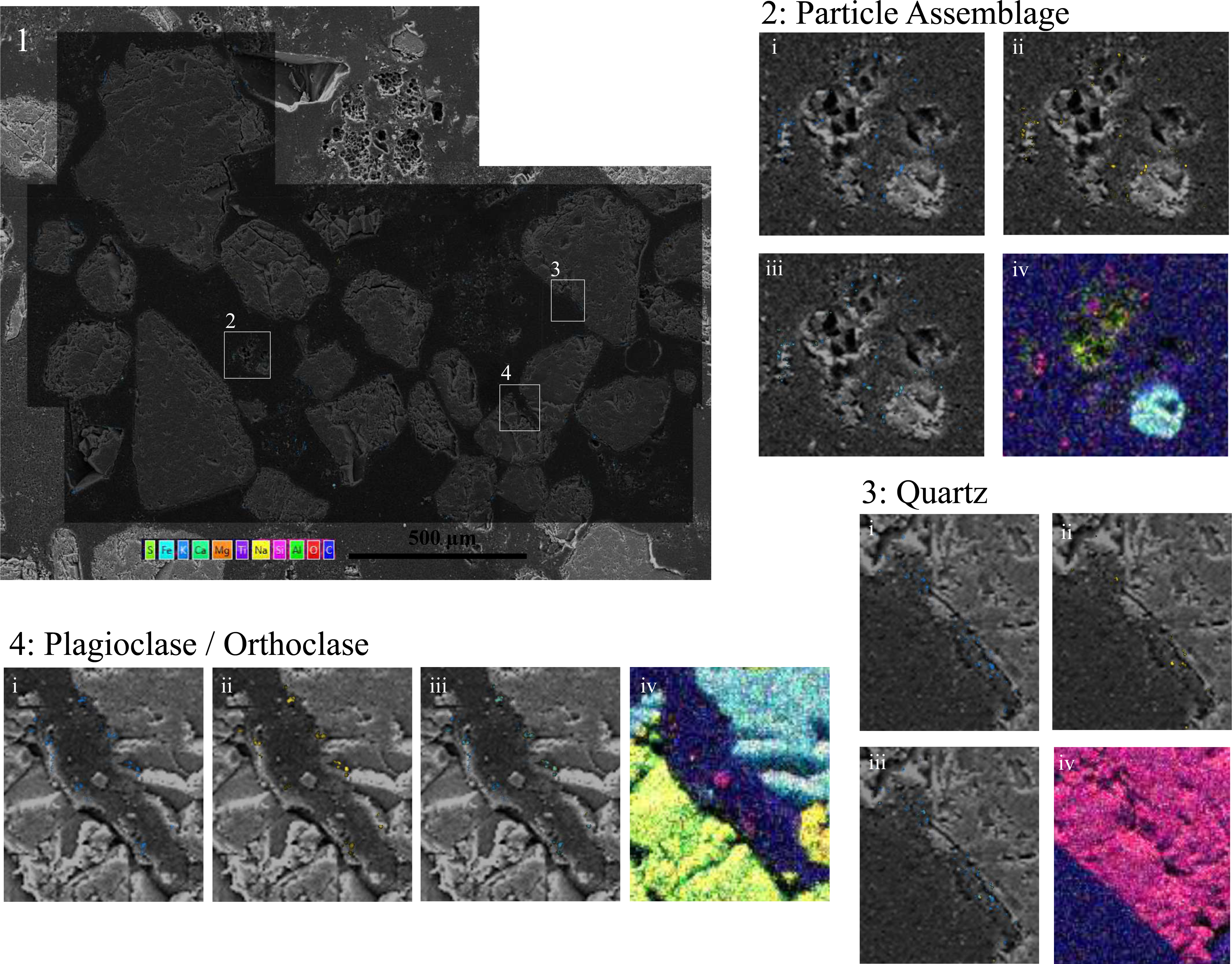
Correlative fluorescence and electron microscopy from the embedded section at 12 mm sediment depth. 1) Overlain on the base SEM image are two fluorescence channels showing SYBR-active features in blue, and BONCAT-active features in yellow. The dark zonation indicates the fluorescence microscopy footprint. 2), 3), and 4) show three mineralogically distinct sites in additional detail in i) SYBR green, ii) BONCAT, and iii) merged channels, as well as iv) EDS elemental abundance maps (in which dark blue background represents the resin).

**Fig 4:**
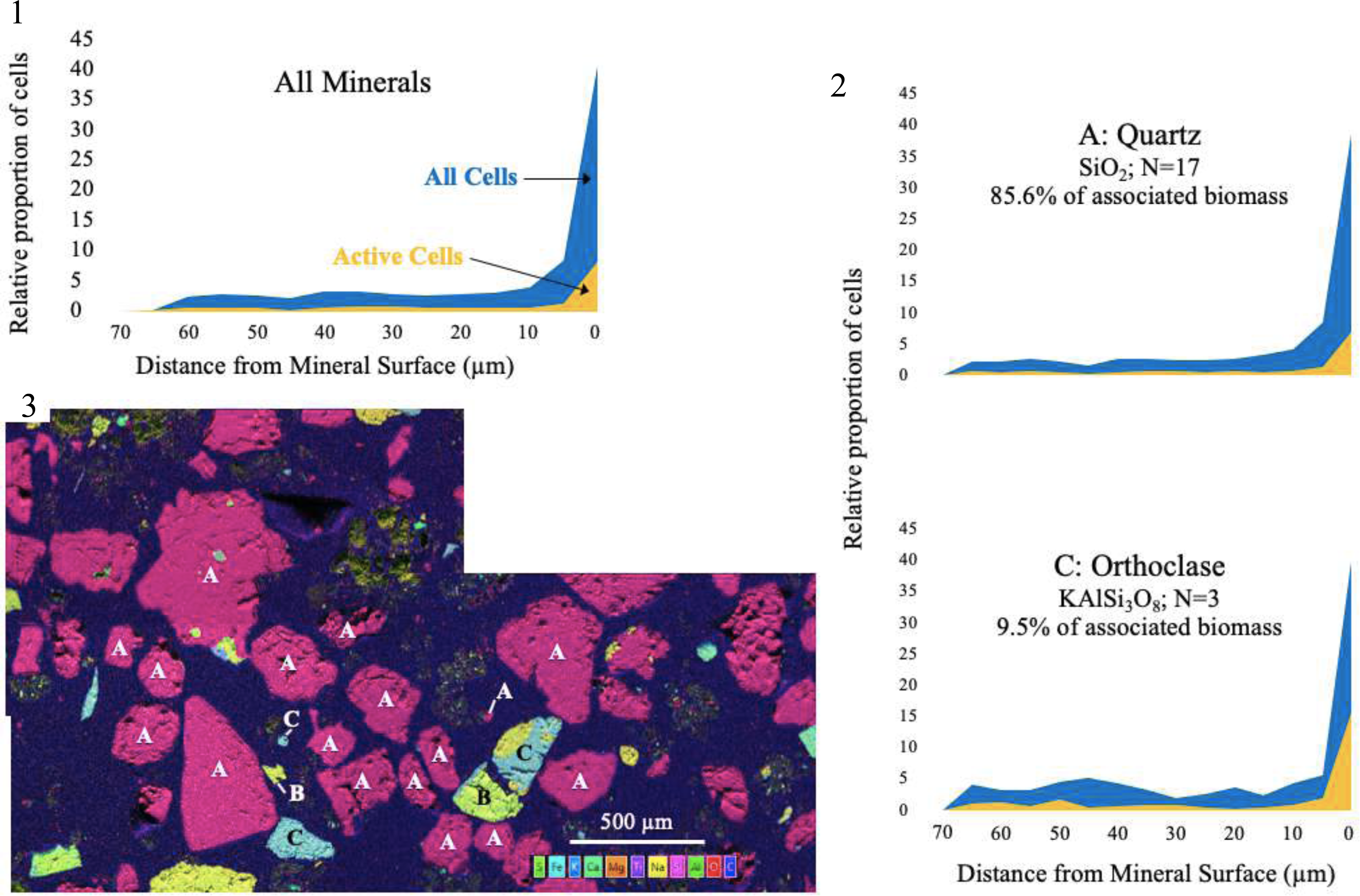
Histograms of the relative proportions of all organisms and the anabolically active subset (yellow overlay) at given distances from the mineral surface for the middle analyzed section (12 mm sediment depth). 1) Data for all grains. 2) Data separated by mineral type. Histogram bins are in 5 µm intervals, and only cells located outside mineral surfaces are shown. 3) Composite elemental maps derived from EDS analysis show the mineral grains that were analyzed, labeled by mineral type. A = quartz; B = Plagioclase; C = Orthoclase; D = Rutile; E = Albite; F = Ca,K,Mg,Fe silicate; G = Hornblende.

### Lower horizons BONCAT-FACS

The horizons from 20-60 mm did not have any corresponding sections analyzed by microscopy, but the FACS and 16S rRNA gene sequencing data illuminate important trends in community composition and metabolic activity with sediment depth. Among the eight most abundant orders recovered, the sulfate-reducing *Desulfobacterales* and *Desulfarculales* accounted for relatively consistent proportions of the active and inactive subsets throughout the core (Fig. 5). The prominence of these lineages is consistent with previous observations that sulfate reduction is the main remineralization metabolism in salt marsh sediments, accounting for roughly 80% of all respiration at Great Sippewissett Marsh (Howarth and Teal, 1979). The more abundant *Desulfobacterales* were more prevalent among anabolically active than inactive organisms at all horizons, while the *Desulfarculales* frequently exhibited the opposite relationship. The latter order consisted of the *Desulfatiglans* genus, whose abundance in subseafloor environments has been attributed to its metabolic versatility in the degradation of aromatics (Jochum *et al*., 2018). In our context, this versatility has seemingly enabled the genus to persist throughout the core, but the cost of a diverse metabolic portfolio could be substantial lag times in metabolic re-routing or extended periods of quiescence for organisms whose metabolic substrate is not present at a given time.

**Fig. 5:**
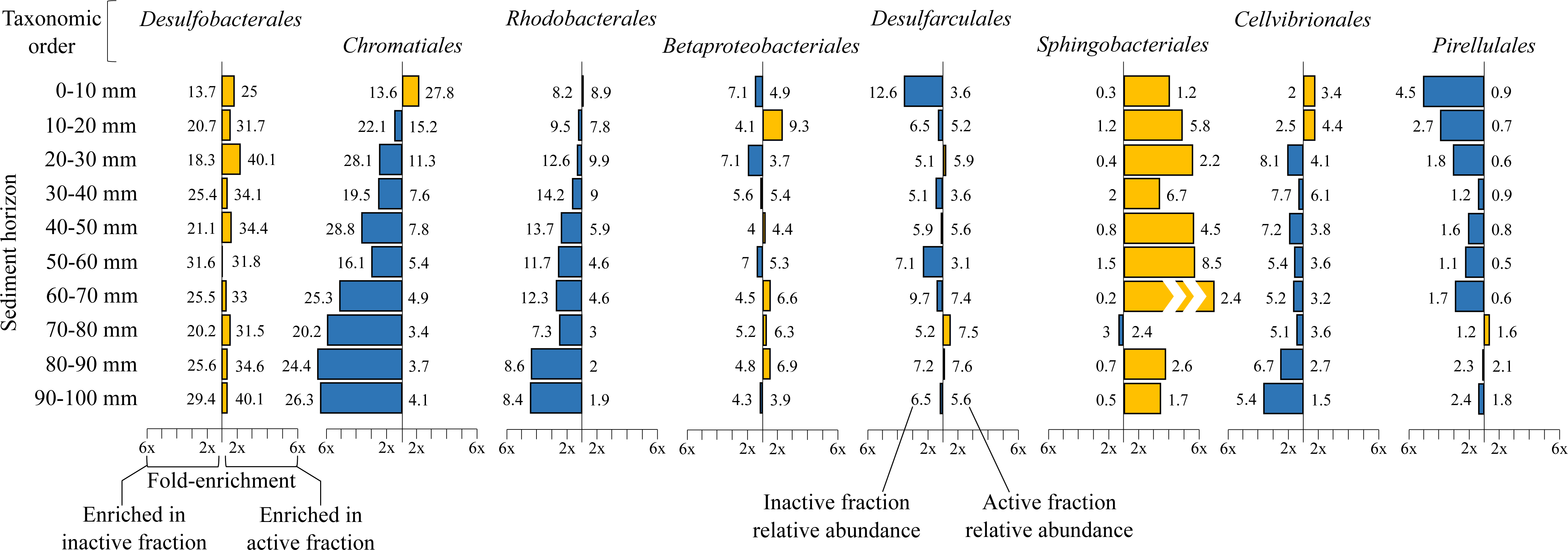
Trends of the relative abundances of active and inactive subsets of the eight most prevalent orders with sediment depth, as detected by BONCAT-FACS combined with 16S rRNA gene sequencing (n=3). At each horizon, the relative abundance contribution for each order was determined in both the anabolically active sorted cells and the inactive sorted cells. Values to the right of the axis indicate the relative abundance of that order in the active fraction; values to the left indicate the relative abundance in the inactive fraction. The colored bars reveal if the order was enriched in the active fraction (yellow bars) or the inactive fraction (blue bars) in a given horizon. The length of bars show fold-enrichment, as indicated by the x-axis, calculated by dividing the larger relative abundance value by the smaller relative abundance value for each horizon.

Purple sulfur bacteria *Chromatiales*, as anticipated, comprised a decreasing proportion of active cells down-core in the absence of light. However, it was the most abundant order in several inactive fractions, suggesting that purple sulfur bacteria may be among the larger microbial contributors of organic matter to deeper sediments. *Cellvibrionales* and *Rhodobacterales* were found at higher relative abundance in active than inactive fractions at the top of the core, but the opposite was true below 20 mm depth. *Cellvibrionales* have traditionally been considered oligotrophs, but some members of the order contain sulfur oxidation pathways and others can grow photoheterotrophically (Spring *et al*., 2014, 2015); this diversity of environments may explain their relatively consistent presence among both active and inactive sequences throughout the core. *Rhodobacterales* are noted early colonizers of particles (Dang *et al*., 2008); one of the most prominent genera detected throughout the core was *Rubribacterium*, a non-sulfur purple bacterium that is a facultative aerobe (Boldareva *et al*., 2009). These traits help explain the order’s presence at all horizons and its decrease in the active fraction with depth.

The observed vertical profile of *Pirellulales* sequences is consistent with aerobic chemoorganotrophs (Schlesner *et al*., 2004) which may have been deposited onto the sediment surface, metabolically inactivated quickly upon burial and the onset of anoxic conditions, and potentially scavenged by the anoxic heterotrophs. *Sphingobacterales* are typically associated with carbon remineralization in oxic soils (Fierer *et al*., 2007), but they do retain fermentation-associated genes (Hester *et al*., 2018) that may explain their presence among the active cell fraction we recovered from below 10 mm depth.

### Deepest horizon BONCAT-FACS & Correlative Microscopy

The deepest section used for correlative microscopy analysis was at a depth of 60.7 mm. Prior to cell extraction and sorting, a diverse range of fermentative lineages was observed, including *Anaerolinaceae* (6.4% of bulk sequences), *Clostridia* (2.6%), and *Bacteroidia* (13%). Few sequences from putative methanogens were observed, potentially due to primer bias (Bahram *et al*., 2019), seasonality (Buckley *et al*., 2008), and the presence of abundant sulfate-reducing bacteria and a range of homoacetogens that may be more successful at attaining hydrogen (Oremland and Polcin, 1982; Ye *et al*., 2014). In this horizon, the majority of observed (87.9%) and sorted (85.5%) cells were anabolically inactive (Table 2). The active and inactive communities exhibited similar richness, but the active community had higher evenness (Fig. S3) and included a comparatively higher relative proportion of *Desulfobacterales*, *Bacteroidia*, *Clostridia*, and *Anaerolinaceae*.

Forty-two mineral grains were observed in the fluorescence microscopy field of view, which also contained the highest abundance of small mineral particles and heterogenous patches of the three sections (Figs. 6-7), potentially due to the diagenetic processes that accompany burial and longer residence times within the sediment column (Curtis, 1987). Despite the high abundance of associated mineral interfaces across a range of spatial scales, this horizon exhibited the lowest microbial abundance and the lowest proportion of anabolically active organisms. This observation is consistent with commonly observed trends in sediments, where electron acceptor depletion and the progressive loss of labile carbon with depth can lead to energetically constrained conditions (Blume *et al*., 2002; Jörgensen *et al*., 2002; Stone *et al*., 2014).

**Fig. 6:**
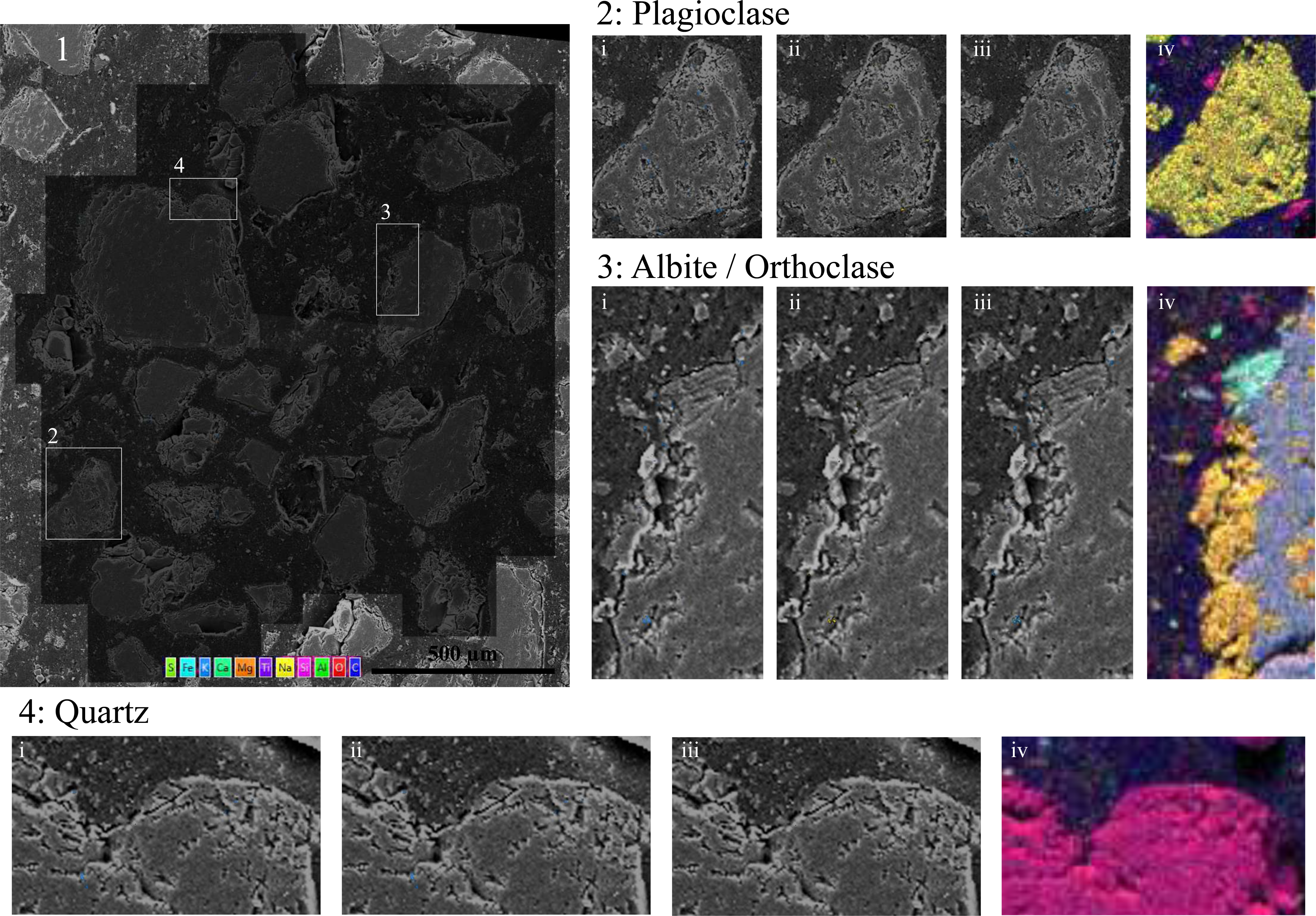
Correlative fluorescence and electron microscopy from the embedded section at 60.7 mm sediment depth. 1) Overlain on the base SEM image are two fluorescence channels showing SYBR-active features in blue, and BONCAT-active features in yellow. The dark zonation indicates the fluorescence microscopy footprint. 2), 3), and 4) show three mineralogically distinct sites in additional detail in i) SYBR green, ii) BONCAT, and iii) merged channels, as well as iv) EDS elemental abundance maps (in which dark blue background represents the resin).

**Fig. 7:**
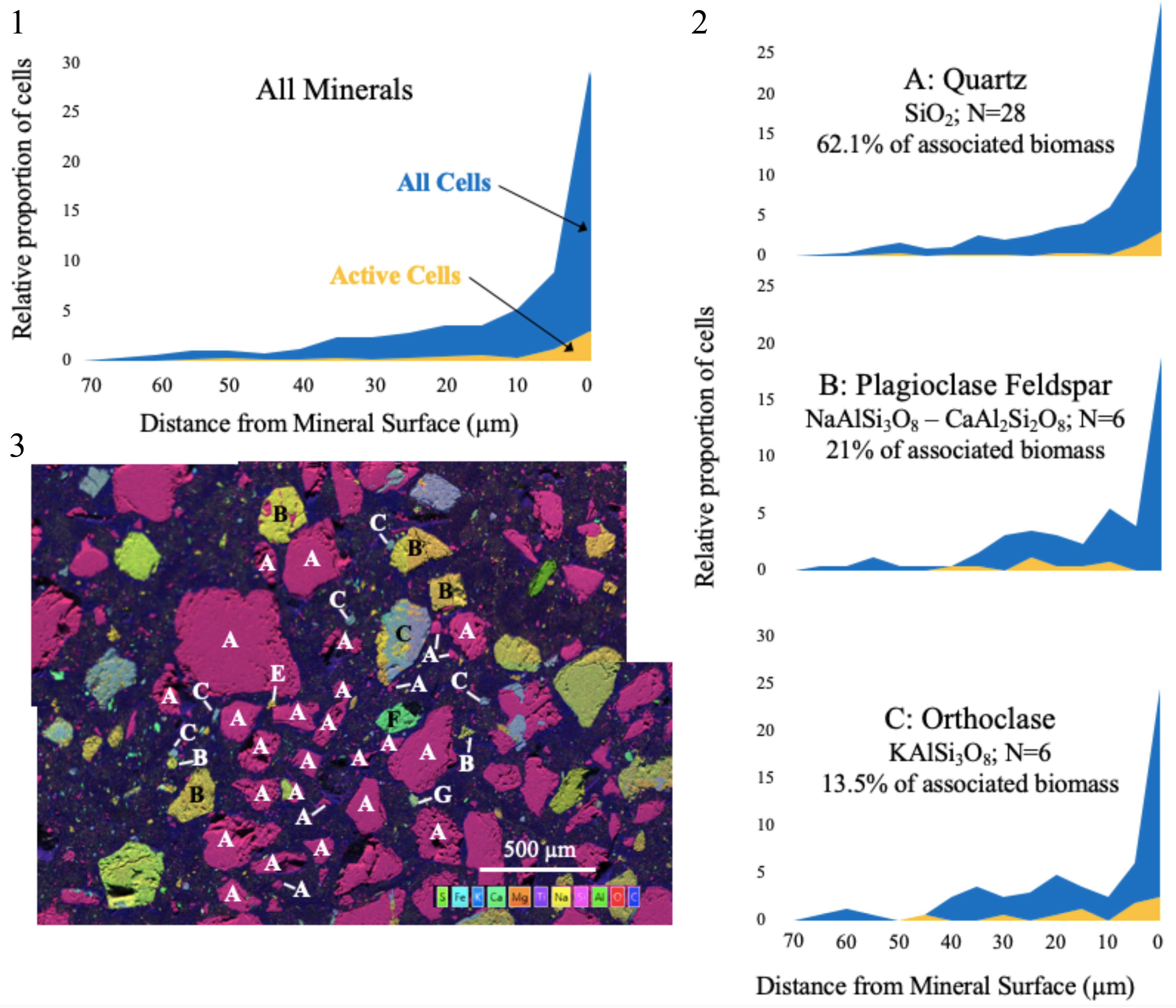
Histograms of the relative proportions of all organisms and the anabolically active subset (yellow overlay) at given distances from the mineral surface for the lowest analyzed section (60.7 mm sediment depth). 1) Data for all grains. 2) Data separated by mineral type. Histogram bins are in 5 µm intervals, and only cells located outside mineral surfaces are shown. 3) Composite elemental maps derived from EDS analysis show the mineral grains that were analyzed, labeled by mineral type. A = quartz; B = Plagioclase; C = Orthoclase; D = Rutile; E = Albite; F = Ca,K,Mg,Fe silicate; G = Hornblende.

In the 60.7 mm horizon, quartz grains had the lowest cell abundances per unit surface area and volume of the three examined sections, while orthoclase and plagioclase had higher-than-average biomass densities (Table 2). The proportion of anabolically active organisms, however, was not substantially different among distinct mineral types, suggesting that cells adhere more strongly to plagioclase and orthoclase grains, and/or that quartz is more readily degraded during diagenesis, disrupting surficial microbial association. This horizon also exhibited the highest proportion of cells located inside mineral grains (37.8%), an observation that could reflect the extensive remineralization of external biomass with burial (Mackin and Swider, 1989).

### Compiling findings across horizons

When integrating sequencing and microscopy data across all horizons, intriguing trends of anabolic activity, diversity, and spatial arrangement emerged. With increasing depth into the sediment, where geochemical and thermal conditions were more stable, alpha diversity metrics of bulk pre-extraction communities revealed a decrease in richness but increase in evenness (Fig. S3). Among the anabolically active and inactive communities, no substantial change in the number of distinct ASVs with depth was observed, but the evenness of their distribution increased down-core for the active constituents. This pattern may reflect a wider range of available niches with fewer dominant lineages below the photic zone, as organic matter is remineralized through a range of metabolic routes, making these deeper communities’ emergent effects more resistant to environmental changes (Wittebolle *et al*., 2009).

Beta diversity analysis revealed a clear separation of active and inactive communities, confirming that organisms respond to environmental cues in a taxonomically differentiated manner and that anabolic activity is not a random process (Fig. S4). Furthermore, for both active and inactive communities, the closer two sediment horizons were in depth, the more similar their community compositions. This trend likely reflects depth-based gradients that form the energetic basis for metabolic activity, as well as the burial process in which a given horizon’s community represents the confluence of local selective pressures operating on an assemblage of organisms “imported” from above or from below due to tidal pumping.

At each of the three horizons examined through correlative microscopy, quartz was the dominant mineral type, yet microbial communities associated with quartz grains had the lowest proportion of anabolically active members. Other mineral types – such as orthoclase, plagioclase, and rutile – had a broader set of cations (Al, Ti, K) that may have offered additional electron transfer or nutrient acquisition opportunities for active cells. With increasing sediment depth, organisms were more likely to be located inside mineral grains, and these “internal” cells were increasingly likely to be anabolically active compared with their “external” counterparts (Table 2): at 7.6 mm, 12 mm, and 60.7 mm depth, internal organisms were 2.1%, 24%, and 45% more likely to be active than those outside minerals. These observations are consistent with a more stable intra-mineral environment that may be less susceptible to predation, particularly in the more energetically constrained anoxic sediment horizons.

## Conclusions

The biological community of LSSM sediment demonstrated notable differences in its composition, anabolic activity patterns, spatial arrangements, and mineralogical associations at the three distinct horizons analyzed in this study. Following incorporation of HPG into new biomass during a 3.7-day *in situ* incubation experiment, correlative microscopy, BONCAT-FACS, and sequencing demonstrated that the most prevalent active constituents shifted from sulfur cycling phototrophic consortia in the surficial horizon, to sulfate-reducing bacteria likely oxidizing a range of organic compounds, to a range of fermentative lineages in the lower horizons. We observed a rapid decay in the proportion of active organisms from ∼60% in the top cm to between 10-25% in the horizons between 2-7 cm depth, offering a quantifiable reflection of the shift to the dark, anoxic environment. By embedding sediment cores in resin, we mapped biomass and mineral grains with microscale resolution and found that, on average, organisms were more distant from mineral grain surfaces in the uppermost horizon, and most likely to be found inside mineral grains in the lowermost horizon. Plagioclase, orthoclase, and rutile minerals recruited more abundant communities that contained a higher proportion of anabolically active organisms compared with quartz grains. Taken together, these findings give the impression of a more spatially and metabolically expansive community in surface sediments, fueled by sunlight and a range of available niches, that is streamlined by burial and mineralogical weathering.

This benchmark study presents a promising new approach for exploring the anabolic activity of a complex microbial community by mapping the precise spatial configuration of anabolically active organisms within mineralogically heterogeneous salt marsh sediment through correlative fluorescence and electron microscopy, while simultaneously identifying active organisms in neighboring sediment with BONCAT-FACS and 16S rRNA gene sequencing. The structure, activity, and evolutionary trajectory of complex microbial communities are determined by the interactions between biotic and abiotic components of an ecosystem. Spatial relationships are a powerful indication of these interactions, particularly in concert with the identification of metabolically active organisms. Looking forward, the incorporation of rRNA-targeted FISH into this workflow would enable a more direct connection between microbe-mineral spatial arrangements and taxonomically constrained activity patterns. Improved approaches for understanding microscale ecosystems in a new light, such as those presented here, reveal environmental parameters that promote or constrain metabolic activity and clarify the impact that microbial communities have on our world.

## Experimental Procedures

### Incubation Chamber Construction

Customized chambers were constructed to enable *in situ* incubation with HPG-infused fluid. Glycol-modified polyethylene terephthalate (PETG) tubes (1” outer diameter, 0.75” inner diameter, McMaster-Carr, Elmhurst, IL) were cut to ∼30 cm length. PETG was used because of its low gas permeability and high optical transparency (<u>Thermo Fisher Scientific</u>), properties that diminished oxygen penetration of subsurface sediments during recovery and transport while retaining light availability for surface-exposed organisms during the incubation period. The lower opening of each tube was beveled with sandpaper (giving the tube a sharp interior edge) to minimize the effects of compaction on collected material when pressing the tube into the sediment.

To make the chambers water-tight, the top portions of two 50-mL Falcon tubes were cut off and attached to either end of the PETG tube using Master Plumber epoxy putty (William H. Harvey, Omaha, NE). By threading the lids onto the appended tube tops, fluid was retained within the incubation tube; by removing them, percolation was enabled.

At certain times during the incubation, gas-permeable, liquid-impermeable conditions were required. This was achieved with 0.01”-thick silicone polydimethylsiloxane (PDMS) membranes (Interstate Specialty Products, Sutton, MA) that were secured between the end of the tube and the screw-top lid. Holes were poked into the Falcon tube lid with a needle to facilitate gas exchange at both the top and bottom of the incubation volume. All materials were thoroughly cleaned with 70% ethanol and 60 minutes of UV light exposure prior to use.

### Sample Recovery, HPG Addition, and *In Situ* Incubation

On September 26^th^, 2018, several customized incubation chamber tubes were taken to Little Sippewissett Salt Marsh (LSSM) in Falmouth, MA. The “Berry Pond” at 41.5758° latitude, −70.6394° longitude (Fig. S1) was selected for sampling due to its extensive heritage in environmental microbiology research, which would provide greater context to our studies. The work presented here pertains to sediment cores collected with five customized incubation tubes (see Table S1). Cores BM and BS were treated with HPG for BONCAT analysis; CM and CS were control samples with no HPG exposure. BM and CM were used for microscopy analysis; BS and CS were used for community analysis by FACS and subsequent 16S rRNA gene sequencing of anabolically active and inactive populations. An additional control of homogenized 0-10 cm depth LSSM sediment (core AM) was autoclave-sterilized and then incubated with HPG solution (described below) for 89 hours in the lab.

At approximately 07:00 (24-hour clock), the tubes were pressed into the sediment, collecting ∼10-12 cm of sediment; upon removal, an autoclave-sterilized plug of glass wool (Fisher Scientific) was inserted into the bottom of the tube. (See Fig. 8 for a schematic of the collection and sample processing approach.) Permeable and intact Falcon tube caps were twisted onto the top and bottom of the chamber, respectively. At approximately 7:30, the incubation chambers were placed in an anoxic glove box (3.5% H_2_, 20% CO_2_, 76.5% N_2_), the caps were removed, and fluid replacement began. Based on the permeability of the sediment and the glass wool plug, fluid moved through the sediment column at a rate of ∼0.3 mL per minute per cm^2^.

**Fig. 8:**
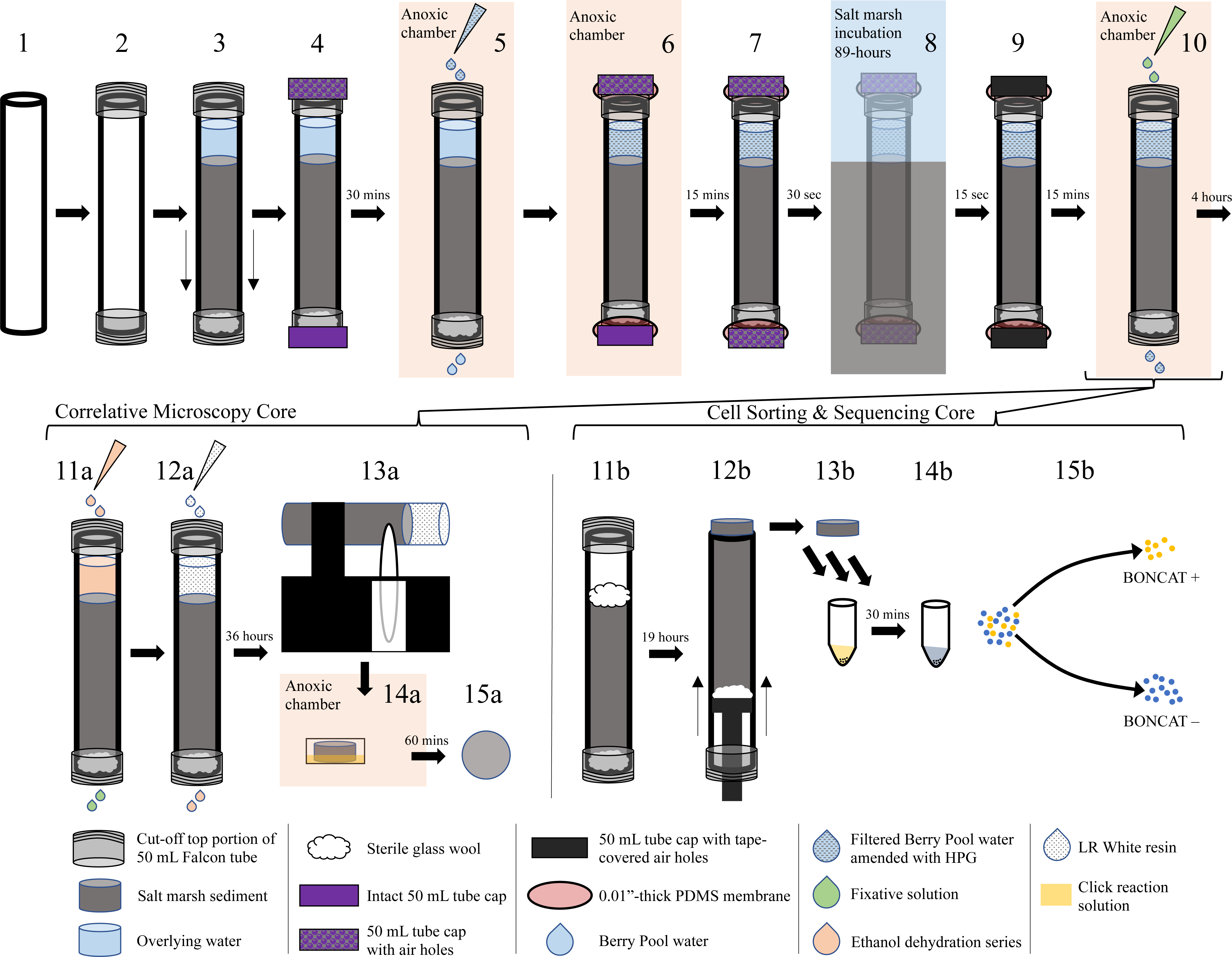
An overview of the experimental and sample processing approach deployed in this study. The PETG tube is cut to the appropriate dimensions and the lower edge is beveled (1). Cut-off 50 mL Falcon tube tops are secured to the PETG tube with epoxy (2), and sediment is collected from the marsh by pressing the tube downward into the sediment (3). A sterile plug of glass wool is added to the bottom to keep the material in place, and the full tube is pulled back out of the marsh. Tube lids are secured; the top lid has a perforated top to allow contact with an oxic atmosphere (4). In an anoxic chamber, lids are removed and fluid is replaced drop-wise by pipette with 50 µM HPG in 0.22 µm-filtered Berry Pool water (5). (Not all cycles of fluid replacement are shown; see text for full protocol.) PDMS membranes are secured to top and bottom of tube with twist-on lids (6). Sample tubes are returned to the marsh; immediately prior to emplacement in the Berry Pool sediment, the bottom lid is perforated to allow gaseous continuity with the environment (7). The sample is placed back in the sediment at the initial collection location for the duration of the incubation period (8); upon recovery, lid perforations are immediately covered with electrical tape to minimize gas exchange during transport back to the lab (9). In the anoxic chamber, incubation fluid is replaced with fixative and incubated for four hours at room temperature (10). Correlative microscopy cores are processed according to steps 11a-15a. The fixed core is removed from the anoxic chamber and infiltrated with an ethanol dehydration series (11a) followed by LR White resin (12a), which is allowed to cure during a 36 hour incubation at 60 °C. The embedded core is then sectioned by sterile water-cooled diamond saw (13a), and sectioned surfaces are incubated in the click solution for 60 minutes in the dark in an anoxic chamber (14a). Sample sections are now ready for SYBR green counterstaining and fluorescence and electron microscopy (15a). Cores for cell sorting and sequencing are processed according to steps 11b-15b. The overlying liquid and top 1.0 cm of sediment is removed and replaced by a plug of sterile glass wool for transport (11b). A sterile plunger was used to extrude the core in 1 cm increments (12b). Cells were extracted from these subsamples and then incubated in the click solution for 30 minutes in the dark (13b). Cells were then washed (14b) and introduced to the cell sorter, which separated BONCAT positive and BONCAT negative cells (15b) for downstream sequencing.

Next, the sediment water was replaced with fluid containing HPG (Click Chemistry Tools, Scottsdale, AZ) for anabolic uptake. 50 µM HPG was dissolved in 0.22 µm-filtered Berry Pond water and stored in an anoxic chamber overnight. (The appropriate concentration of HPG and dye for subsequent visualization was determined from a previous study (Hatzenpichler and Orphan, 2015) and a series of tests using 5-500 µM HPG and 0.5-50 µM dye; see Fig. S5.) Four to six column volumes of HPG solution were added dropwise to the top of the column at roughly the same rate as fluid leaked out the bottom, resulting in a full replacement of permeable volume with HPG solution in a manner that did not substantially change the overlying pressure experienced by the sediment. PDMS membranes were secured to the top and bottom of the incubation chamber with the same Falcon tube caps as before, and the samples were transported back to the Berry Pool.

At the precise location of sample collection, the incubation chambers were placed back into the marsh sediment at 12:00. Immediately prior to deposition, holes were poked in the bottom caps (but not the membrane) to ensure the full length of the sediment was in gaseous equilibrium with its surroundings, but that the HPG solution remained contained. The tubes were aligned such that the water-sediment interface matched that of the sediment in the Berry Pool. The samples were left to incubate in the marsh for 89 hours (retrieval time September 30^th^, 05:00). The incubation timing was determined by experiments of LSSM sediments demonstrating apparent saturation of BONCAT-positive signal after 88 hours (Fig. S6).

Throughout the process described above, the incubation chambers were checked for leaks. Overlying water levels were marked on the tube exterior after sample recovery, after HPG fluid introduction, and upon re-introduction to the marsh; experiments only proceeded if no change in water level was observed.

### Preparation of Cores for Microscopy

Our core preparation expanded on a protocol that was first used in the analysis of volcanic fumarole soils (Marlow *et al*., 2020). Cores BM and CM were chemically fixed, dehydrated, embedded in resin, sectioned, and stained for analysis by fluorescence and electron microscopy. Four to six column volumes of 3% PFA solution (in 0.22 µm-filtered Berry Pond water) were percolated through the samples to fully replace the HPG solution. An intact bottom cap was secured to the bottom of the core tube, and samples were incubated at room temperature for four hours. A dehydration series consisting of 4-6 column volumes each of 50%, 80%, and 96% ethanol (in 0.22 µm-filtered Berry Pond water) was performed.

Following dehydration, sediment samples were embedded in LR White resin (hard grade, Electron Microscopy Sciences, Hatfield, PA), which was selected for its low viscosity and minimal background signal under the wavelengths used for fluorescence microscopy. LR White has been used in a number of similar applications, including correlative microscopy of animal tissue (Hegermann *et al*., 2019), plant tissues (Bell *et al*., 2013), low diversity microbial biofilms (Knierim *et al*., 2012), carbonate microbialites (Gérard *et al*., 2013), and marine sediment (McGlynn *et al*., 2018). Four to six column volumes of 100% liquid resin were percolated through the dehydrated columns, and intact Falcon tube caps were secured on the bottom to avoid leakage. The samples were then placed in an incubation oven at 60 °C for 36 hours to cure.

The solidified sediment columns were sectioned with a diamond saw (Model 650, South Bay Technology, San Clemente, CA) and an ethanol-sterilized PELCO diamond wafering blade (#812-332, Ted Pella, Inc., Redding, CA) spun through ultrapure Milli-Q cooling water. Sections were submerged in a 5 µM Cy3 Picolyl Azide dye (Click Chemistry Tools, Scottsdale, AZ) click staining solution (Hatzenpichler and Orphan, 2015) in an anoxic chamber for 60 minutes in the dark. Afterwards, they were removed from the chamber, washed three times with sterile PBS solution, incubated in 5x SYBR Green I (referred to hereafter as “SYBR green”; Life Technologies, ThermoFisher, Waltham, MA) in the dark at room temperature for 15 minutes, rinsed three times with sterile PBS, and left to air dry in the dark prior to imaging. All downstream correlative microscopy was performed on areas as far from the outer edge of the sediment core as possible (∼8 mm) in order to minimize the effects of the coring process on the analyzed area.

### Fluorescence Microscopy

Fluorescence imaging of sectioned samples was done with a LSM 880 confocal laser scanning microscope (Zeiss, Oberkochen, Germany) equipped with a gallium arsenide phosphide (GaAsP) detector, a 20x objective lens, and DI water immersion. Argon and DPSS lasers provided excitation at 458, 488, 514, and 561 nm wavelengths. Detected emission windows were 510-561 nm for SYBR green and 564-669 nm for Cy5. Reflected light from the 488 nm laser was also captured to link sample features between confocal and electron microscopy images as reference points. Imaging was done with the Zen 2.3 SP1 program (Zeiss, Oberkochen, Germany). Focus was adjusted manually for each field of view. 1,024 x 1,024 pixel frames were acquired with a pixel dwell time of 32.77 µsec and a pixel size of 240 nm. Four line-based scans were averaged to generate each image. Gain settings were set to minimize background and nonspecific signals. For the SYBR channel, 800 master gain and 1.3 digital gain were used; these parameters were 680 and 1.2, respectively, for the Cy3 channel.

### Electron Microscopy & Energy Dispersive X-Ray Spectroscopy

After fluorescence microscopy, each section was dipped in a dehydration series of 50%, 80%, and 100% ethanol solutions (balance Milli-Q water). The sample was then mounted on a SEM sample holder with double-sided carbon tape and sputter-coated with 5 nm of Pt/Pd (EMS 150 T S Metal Sputter Coater, Electron Microscopy Science, Hatfield, PA). Using the Zeiss SmartSEM software, secondary electron images were collected at a voltage of 12 kV using an Everhart-Thornley detector. The voltage was increased to 20 kV for elemental analysis. A silicon drift detector was used, and data were processed with the EDAX Genesis software (Ametek, Berwyn, PA). Elemental maps were used for visualization purposes, supported by quantitative area-based scans to inform mineral identification by X-Ray diffraction.

### X-Ray Diffraction

X-ray diffraction (XRD) was performed using an X’Pert3 powder diffractometer by Panalytical using a Cu Kα source to scan from 5-70**°** 2θ. The sample was prepared as a packed powder and was scanned wobbling the sample stage at 0, −1, and +1 degrees. The final scan was an average of these three scans. Phase identification and semi-quantitative analysis of diffractograms was performed using the HighScore Plus software by Panalytical (Malvern Panalytical, Malvern, United Kingdom).

### Image Processing

Fluorescence microscopy images were processed in Fiji / ImageJ version 2.0.0-rc-69/1.52p (Schindelin *et al*., 2012). We used the DeconvolutionLab2 plugin with a point spread function calculated by the Zen program (based on our specific imaging parameters) and five iterations of the Richardson-Lucy algorithm (Richardson, 1972; Lucy, 1974). The Despeckle program was used for denoising.

To link the location of fluorescent signal with the high-resolution textural information enabled by electron microscopy, image co-registration was performed using the bUnwarpJ algorithm using the following parameters: accurate registration, very coarse initial deformation, super fine final deformation, 0.1 divergence weight, 0.1 curl weight, 3.5 landmark weight, 0.0 image weight, 10 consistency weight, and 0.1 stop threshold. The SEM/EDS images were designated as the “target” images onto which fluorescence images were mapped, and approximately one landmark per 10,000 µm^2^ was designated on the SEM and reflected light fluorescence images. The other two fluorescence channels (SYBR green and Cy3) were anchored to the reflected light layer to accurately co-register SYBR and BONCAT signals with SEM and EDS data (see Fig. S7).

Cell counting and distance relationships were analyzed in Matlab R2018b. Red (Cy3, BONCAT) and green (SYBR green, all cells) channels were separated and converted to binary images at a manually determined global threshold of 0.04. Single pixels were eliminated to remove noise, and individual regions of interest were designated by applying a watershed transform with four-degree connectivity (*e.g.*, pixels that only touched at corners rather than edges were not counted as the same object). Finally, centroid holes were filled, and remaining shapes were counted as “organisms”. An organism was recorded as anabolically active if at least one of its pixels fluoresced both red and green after being subjected to the image analysis pipeline.

Organism concentrations were calculated by counting the number of organisms in the relevant field of view and dividing that number by the volume sampled through fluorescence microscopy. This volume was determined using the x and y dimensions of the microscopy footprint and the z dimension (1.66 µm) empirically established by an average of the SYBR green and Cy3 signal transmission distances through the resin, as shown in Fig. S8.

Distances between organisms and mineral surfaces were measured by first manually tracing the outlines of mineral grains in Adobe Photoshop using the high-resolution SEM images and converting the resulting image into a binary image in Matlab. Next, we calculated the shortest Euclidean distance from each organism’s outer surface to the perimeter of each mineral. The shortest distance and identity of the associated mineral were recorded. Because this analysis was restricted to a two-dimensional cross-sectional view of three-dimensional mineral grains, distances from mineral grain-internal cells to the mineral surface are not reported. To exclude cells that had been dislodged during the embedding and sectioning process, all cells more than 70 µm away from the nearest mineral surface were omitted from distance-based calculations. This cutoff, which removed 6.2% of all organisms from spatial analysis only, was based on a distance histogram to determine outliers (Fig. S9) and is within the range of biofilm thicknesses associated with silica mineral surfaces (which represented the majority of observed mineral grains) (Ye *et al*., 2015). During sectioning, a few mineral grains were plucked from the resin: three in the top section, two in the middle, and three in the bottom. Organisms associated with these grains that remained in the resin were not included in spatial analyses, and the overall loss of material indicates that cell abundance values represent a lower bound. When normalizing organism abundances by mineral surface area and volume, perimeters calculated from mineral grain outlines were used as a proxy for surface area, and cross-sectional area was used as a proxy for grain volume.

### Fluorescence-Activated Cell Sorting (FACS)

Samples BS and CS were used for FACS and high-throughput 16S rRNA gene sequencing to identify the subset of microorganisms that was anabolically active during the incubation period. These sediment cores were shipped on ice from Massachusetts to Montana State University for analysis (shipment took ∼19 hours). Prior to shipment, the overlying liquid was removed and the top 1.0 cm (+/− 0.3 cm) flocculent layer of sediment was transferred to a separate sterile tube to avoid compression and inaccurate identification of horizons. Sterile glass wool was added to the top of the incubation chamber to maintain core coherence during transport.

Upon arrival, each sediment core was carefully excised from the core sleeve using a custom-built, sterilized plunger. Each core was divided into 1 cm increments, which were weighed, transferred into Falcon tubes containing 10 mL of sterilized 1xPBS, and stored at 4°C until cells were extracted. Cells were extracted from each sediment layer using methods adapted from Couradeau et al. (Couradeau *et al*., 2019) with the following modifications. For each sediment layer, 1 mL of the slurry was diluted with 5 mL of sterile PBS in a 15 mL Falcon tube with Tween20 (final concentration 0.02%). The cell extraction slurry was placed on a benchtop vortexer at maximum speed for 5 minutes. Large sediment particles were pelleted via centrifugation at 500 x g for 55 minutes. Cells from 700 µL of the supernatant were pelleted in a 1.5 mL microcentrifuge tube by centrifugation at 16,000 x g for 5 minutes. The supernatant was carefully removed by pipette before the click reaction was performed directly on the cell pellet. Extraction blanks were performed without any added sediment in parallel with cell extractions to test for reagent contamination.

The click reaction solution was prepared in a large volume in order to stain all samples using the same solution. The reaction solution was prepared as previously described (Hatzenpichler and Orphan, 2015) with the addition of a general DNA stain, SYBR green (ThermoFisher Scientific, Invitrogen, Eugene, OR, USA) to counterstain all cells. The solution contained 5 mM aminoguanidine hydrochloride (Sigma Aldrich), 5 mM sodium *L*-ascorbate (Sigma Aldrich), 100 µM copper sulfate pentahydrate (Sigma Aldrich), 500 µM THPTA (Click Chemistry Tools), 12 µM Cy5 picolyl-azide dye (Click Chemistry Tools, Scottsdale, AZ), and 1x SYBR green in PBS. The solution was vortexed, and 200 µL was transferred to each cell pellet and mixed with cells by pipetting up and down. The click reaction solution and cells were incubated for 30 min in the dark on a slow rotator at room temperature. Click reaction solution was washed from cells by three cycles of (1) centrifugation at 17,000 x g for 5 minutes, (2) removal of supernatant by pipette, and (3) resuspension in sterile PBS. The final cell pellet was resuspended in 700 µL of sterile 1x PBS. Prior to cell sorting, the cell suspension was passed through a 35 µm filter cap (BD-falcon 5 mL tube with cell strainer cap, CorningTM, Corning, NY, USA) to remove any remaining large debris.

A Sony SH800S FACS (Sony Biotechnology, San Jose, CA) was used to sort cells via a 70 µm chip. The cell sorter was set to detect the SYBR green dye on the green channel (excitation 488 nm) and Cy5 dye on the red channel (excitation 638 nm). The first two gates were drawn on forward scatter and back scatter properties to remove any large particles or noise. The third gate was drawn on SYBR green positive (“sortable cells”) events, with the assumption that this gate captured all cells. Only SYBR+ cells were gated to be either BONCAT positive (“active cells”) or BONCAT negative (“inactive cells”) on the Cy5 channel. Core CS, the no-HPG control, was used to determine the cutoff between BONCAT positive and negative fractions. Gates were drawn conservatively to minimize the possibility for false positives or false negatives in any gate (Fig. S2). For each of three biological replicates for each cell fraction (“active cells” and “inactive cells”), 1×10^6^ cells were sorted into 1.5 mL microcentrifuge tubes containing 500 µL of sterile 1x PBS. Sorted cells were stored at 4 °C for up to six hours before being centrifuged at 17,000 x g for 5 minutes to pellet and then resuspended in 20 µL of nuclease free water and frozen at −80°C until DNA extraction and downstream processing.

### 16S rRNA Gene Amplicon Sequencing

To capture the bulk microbial community from each sediment layer in core BS, DNA was extracted from 500 µL sediment/1x PBS slurry from each layer using the FastDNA Spin Kit for Soils (MP Biomedicals, Irvine, CA) following the manufacturer’s instructions. A blank DNA extraction was processed in parallel with bulk sediment extractions to check for contaminants. DNA was extracted from sorted cells and processing blanks as previously described (Reichart *et al*., 2020). Briefly, cell suspensions were transferred to a 96-well microtiter plate and sealed with sterile adhesive foil before being subjected to three freeze-thaw cycles (−80 °C for 20 min, 99 °C for 10 min), with brief centrifugation prior to each freezing step. 16S rRNA genes of bacteria and archaea were amplified following the Earth Microbiome protocol (Thompson *et al*., 2017) using revised primers 515F (Parada *et al*., 2016) and 806R (Apprill *et al*., 2015) added directly into the extracted DNA microtiter plates. These primers were designed prior to the discovery of several new lineages related to the DPANN and Asgard archaea superphyla (Baker *et al*., 2020) and thus do not capture the entire archaeal diversity (Bahram *et al*., 2019; Pausan *et al*., 2019). However, their extensive use in environmental microbiology laboratories around the world enables comparability between studies (Gilbert *et al*., 2014; Thompson *et al*., 2017) and within our own laboratories. The final PCR volume was 37.5 µL and consisted of 15 µL Invitrogen Platinum Taq II 2X Master Mix, 0.75 µL 515F primer (10 µM; final: 0.2 µM), 0.75 µL 806R primer (10 µM; final: 0.2 µM), and 1 µL nuclease-free water added directly into the microtiter plates containing the 20 µL of lysate. The thermocycler conditions were: 94 °C for 3 minutes followed by 28 cycles of 94 °C for 45 sec, 50 °C for 60 sec, and 72 °C for 90 sec before a final elongation step at 72 °C for 10 minutes. A negative PCR control was processed in parallel using nuclease-free water instead of extracted DNA to check for PCR contaminants. PCR products were purified using AMPure XR beads (Beckman Coulter) following the manufacturer’s protocol with a final elution in 40 µL nuclease free water.

Afterwards, a second PCR to attach barcodes and sequencing adapters to the 16S rRNA gene amplicons was performed. The final volume of the PCR was 25 µL and contained 5 µL purified, amplified DNA, 12 µL Invitrogen Platinum Taq II 2X Master Mix, 2.5 µL i5 primer (final: 0.25 µM), 2.5 µL i7 primer (final: 0.25 µM), and 2.5 µL water. The thermocycler conditions were 95 °C for 3 minutes followed by 8 cycles of 95 °C for 30 sec, 55 °C for 30 sec, and 72 °C for 30 sec, followed by a final elongation step at 72 °C for 5 minutes. A PCR product purification was performed as described above using AMPure XR beads. Finally, purified PCR products were checked for appropriate length by gel electrophoresis in 1% agarose. Amplicons were quantified in triplicate using Quant-iT Picogreen dsDNA Assay (Invitrogen) following the manufacturer’s protocol and measured on a Biotek Synergy H1 Hybrid microplate reader. Samples were pooled at 10 ng DNA each, and the final pooled sample was purified and concentrated with the QIAquick PCR purification spin column kit (Qiagen) following the manufacturer’s guidelines. Sequencing was performed by Laragen Inc. (Culver City, CA) using Illumina 2×250 paired end read MiSeq sequencing. Sequences have been archived at NCBI Genbank under the Bioproject ID PRJNA643437.

### Sequence analysis

Primers were removed from demultiplexed sequences using *cutadapt* (Martin, 2011) with a max mismatch of 2 bp and requirement that primers must be present on both forward and reverse reads to maintain the read pair in the dataset. Unpaired primer-free reads were processed in *DADA2* (Callahan *et al*., 2016) run in the R environment. Reads were first trimmed to 220 bp for forward reads and 170 bp for reverse reads, then were filtered with default settings of maxN=0, maxEE=c(2,2), trunc=2, and rm.phix=TRUE. Denoising of reads was conducted with *DADA2*’s error model calculated on randomly sampled reads from the entire dataset. Forward and reverse reads were merged with a 20 bp minimum overlap and no mismatches. Chimeras were removed within *DADA2* using the “consensus” method. Taxonomy of the amplicon sequence variants (ASVs) was assigned using the SILVA132 database (Quast *et al*., 2012). To remove contaminating sequences, R package *decontam* (Davis *et al*., 2018) was implemented using the “prevalence” model with a threshold of 0.7, which removed 150 of the total 11,014,619 ASVs. Five samples with fewer than 10,000 sequences were removed from the dataset. Further normalization of read count per sample was performed using the R package *metagenomeSeq* (Paulson *et al*., 2013), which builds a statistical model to account for undersampling. Diversity metrics including Shannon’s diversity index, Bray-Curtis similarity, and non-metric multidimensional scaling (NMDS) were calculated in the *Phyloseq* (McMurdie and Holmes, 2013) R package.

### Quality Control

Analysis of samples subjected to the full experimental treatment alongside control samples allowed us to validate our procedures. Comparing sample BM with AM indicated that neither SYBR nor Cy3 signals were attributable to background fluorescence or non-specific binding of HPG or the dyes; across five fields of view from both samples, 96.6% of SYBR-active objects and 97.1% of Cy3-active objects were present in sample BM. Comparing sample BM with CM revealed that HPG did not affect SYBR signal but was required for BONCAT signal: 44.8% of SYBR-active objects and 97.2% of Cy3-active objects were found in sample BM.

Clarifying the role that our experimental treatment had on the microbial community and the empirical biases that may result was a key priority. Daily fluctuations of the Berry Pool water level, which ranges from ∼5-30 cm water depth over the course of a tidal cycle, consistently introduce and remove transient organisms that may not be physically associated with sediment particles. Nonetheless, it is possible that the percolation of fluids through the incubation chambers might transport microbial cells outside of their naturally-occurring habitats. To test this possibility, we introduced 1 mL of 1×10^9^ / mL 1 µm diameter YG carboxylate fluorescent microspheres (Polysciences, Warrington, PA) to the overlying water of a LSSM sediment core. These microspheres are commonly used to simulate microorganism transport and constrain contamination in sediments, soils, and subsurface environments (House *et al*.; Smith *et al*., 2000; Goeppert and Goldscheider, 2011; Bang-Andreasen *et al*., 2017; Labonté *et al*., 2017; Daly *et al*., 2018). Following microsphere addition, the core was treated identically to the BM sample. Flow-through liquid fractions were collected and deposited on 0.22 µm polycarbonate filters, and multiple horizons were sectioned and examined with fluorescence microscopy. Bead counts over five representative fields of view at 10.7 mm above the sediment-water interface, 2.0 mm depth, 5.3 mm depth, 9.8 mm depth, and 23.3 mm depth (Fig. S10), as well as 16 liquid fractions, were averaged and scaled by the overall cross-sectional area of the core. Z-axis transmission of bead fluorescence under confocal microscopy examination was 8.75 µm. Linear interpolation of data points indicated that 99.3% of beads remained above the 7.6 mm horizon, which was the shallowest horizon used for microscopy analysis. Assuming a cell density of 10^6^ / mL in the overlying water and 30 mL of overlying water in the initial core sample, we calculate that 6×10^−3^ % and 8×10^−4^ % of cells detected in the 7.6 mm and 12 mm horizons, respectively, are attributable to entrained surface water cells. Because a horizon lower than 60.7 mm was not examined with the bead test, an analogous figure is not attainable for the 60.7 mm horizon. However, given the trends observed here, we believe the contribution from surface-entrained organisms is negligible. This analysis gave us confidence in interpreting mineral-associated organisms as native to the observed sediment horizons.

## Acknowledgements

This work was supported through a grant by the Gordon and Betty Moore Foundation (GBMF5999). We thank Dr. Julie Huber, Dr. Mak Saito, and Dr. Jaclyn Saunders for graciously providing lab space in Woods Hole in close proximity to the field site, Dr. Ana Pantelic for fieldwork assistance, and Dr. Amy Gartman for assistance with X-ray diffraction analyses. Some analyses presented here were conducted at the Harvard University Center for Nanoscale Systems (CNS), a member of the National Nanotechnology Coordinated Infrastructure Network (NNCI), which is supported by the National Science Foundation under NSF ECCS award no. 1541959. We thank the Harvard Center for Biological Imaging for infrastructure and support. The authors declare no conflict of interest.

## Supplemental Figure and Table Captions

Fig. S1: Sampling site at Little Sippewissett salt marsh. 1) A satellite image of the marsh acquired on October 6^th^, 2018 (Google Earth). The white box indicates the “Berry Pool” shown in image 2 where the sampling was conducted. The white arrow indicates the direction in which image 2 was acquired. 2) The “Berry Pool”, so named because of its high abundance of phototrophic pink berries, as seen on August 25^th^, 2018. The white arrow points to the site of sample acquisition on September 26^th^, 2018. 3) A closer view of the sediment surface at low tide on September 26^th^, showing pink berries, organic surface cover, and sandy sediment. 4) Custom-built sample chambers placed at the site of collection for incubation.

Fig. S2: FACS plots. (A) Shows the no HPG controls used to draw gates around the BONCAT positive (active) and BONCAT negative (inactive) cell fractions in the HPG-added sediment core (B). Note that the biomass extracted from the no HPG control (A) was much lower than seen in other samples where HPG was added.

Fig. S3: Alpha diversity metrics of the bulk, BONCAT+ (Cy5+), and BONCAT− (Cy5−) communities analyzed by 16S rRNA gene amplicon sequencing for each sediment horizon.

Fig. S4: Beta diversity metrics derived from 16S rRNA gene amplicon sequence data. 1) NMDS comparing bulk community with active/inactive sorted communities (stress 0.0784916). 2) NMDS showing differences in sorted active/inactive communities by depth (stress 0.1517008).

Fig. S5: Representative fields of view of *E. coli* cultures exposed to different concentrations of HPG and the azide dye. The 50 µM HPG, 5 µM dye combination provided the best combination of coverage and dynamic range and was used in the field-based experimental incubation. All samples were stained with the general DNA stain DAPI.

Fig. S6: The percentage of Sippewissett biomass that was anabolically active as a function of incubation time. All incubations used homogenized LSSM sediment from the 0-5 cm horizon, and received 50 µM HPG. Active and inactive organisms were quantified as described in the text. Data points represent mean and standard deviation values across 5 fields of view; one field of view is provided in both SYBR green and BONCAT channels for 1, 44, and 88 hours below the graph.

Fig. S7: To co-register fluorescence and electron microscopy images and facilitate precise spatial analysis, the bUnwarpJ algorithm in FIJI / ImageJ was used. See the text for details on parameter settings.

Fig. S8: To determine the z-axis depth into the embedded section that our protocol would detect, Cy3 (BONCAT) and SYBR green (all cells) channels were recorded at multiple focal depths with a step size of 0.35 µm. Three BONCAT and two SYBR green features are highlighted. Each resulting image was processed identically, as described in the text; Cy3 signal was color-shifted to yellow and SYBR green signal was color-shifted to blue for viewing ease. Features that registered as an “object” after processing have a yellow or blue border; those that were not have no border. For each feature, the depth-based analysis began when the object was fully in focus.

Fig. S9: Histograms of the number of organisms as a function of distance from mineral surfaces. Histogram bins are in increments of 5 µm; the number of each bin corresponds to the upper bound of the range (e.g., “5” includes all organisms between 0 and 5 µm from the mineral surface). Only organisms on the outside of mineral grains are shown. Mineral types with less than 83 associated cells (e.g., <1% of the total mineral-external cells observed in this study) are not shown.

Fig. S10: To test the effect of our fluid replacement approach on transport of microbial cells, 1-micron fluorescent beads were introduced to the overlying water and tracked through the core during the fluid replacement and embedding process. Horizons of fluorescent bead quantification are indicated by green arrows, and representative fields of view are shown at right. Horizons analyzed for correlative microscopy and FACS-16S rRNA gene sequencing are shown with black arrows and blue brackets, respectively.

Table S1: Details on the conditions and analyses to which experimental and control sediment cores were subjected.

