## Supplementary figures and images for "Spatially-resolved correlative microscopy and microbial identification reveals dynamic depth- and mineral-dependent anabolic activity in salt marsh sediment"

### Figure S1

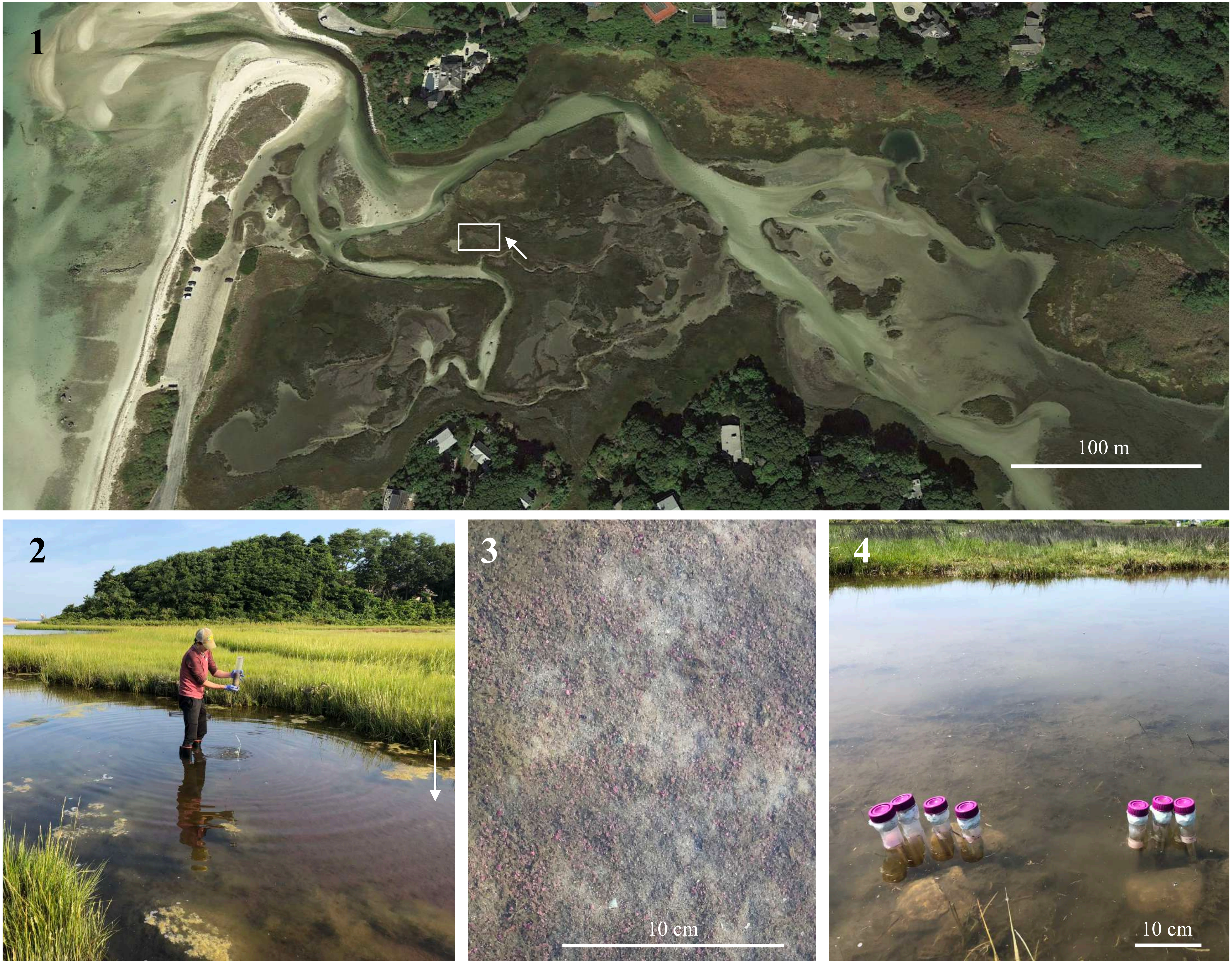

### Figure S2

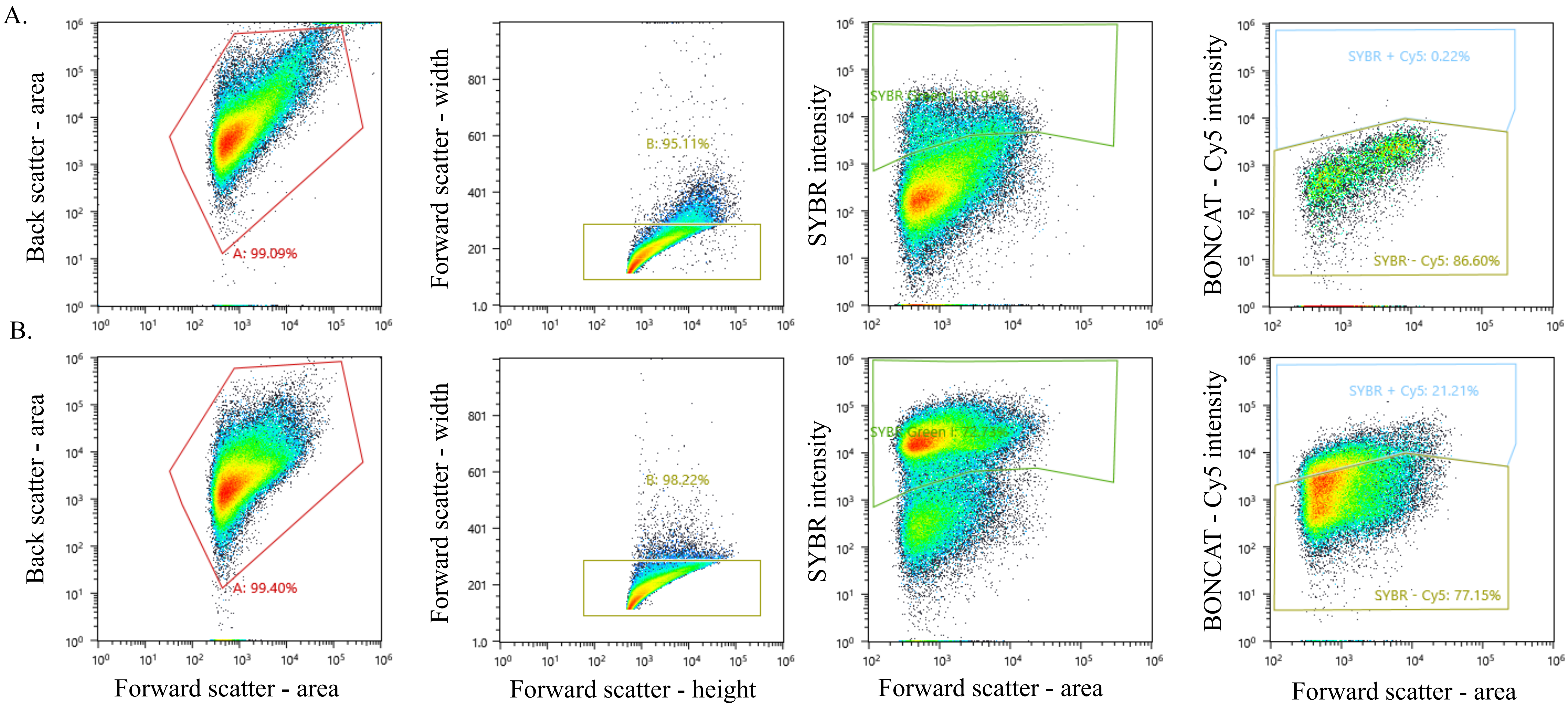

### Figure S3

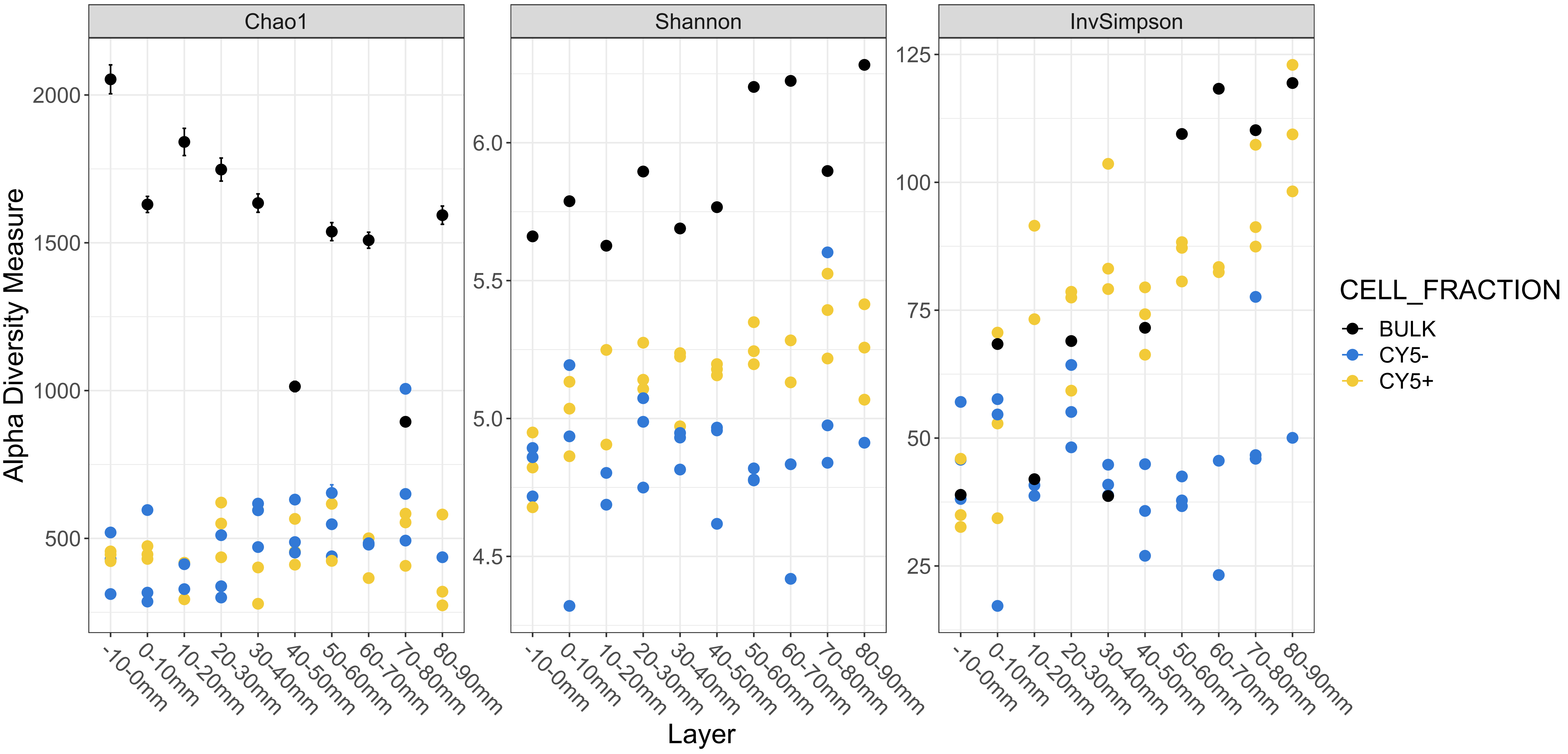

### Figure S4

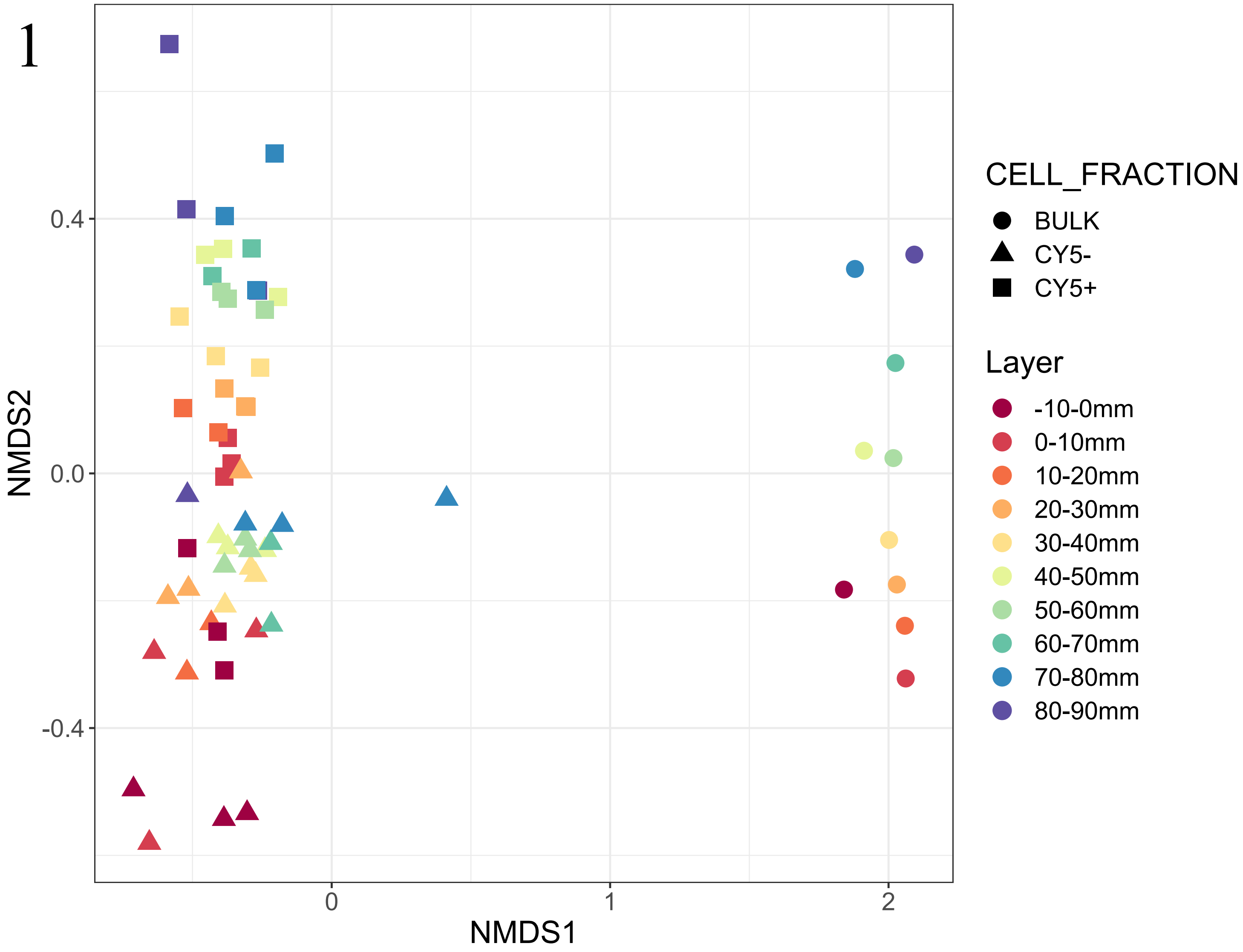

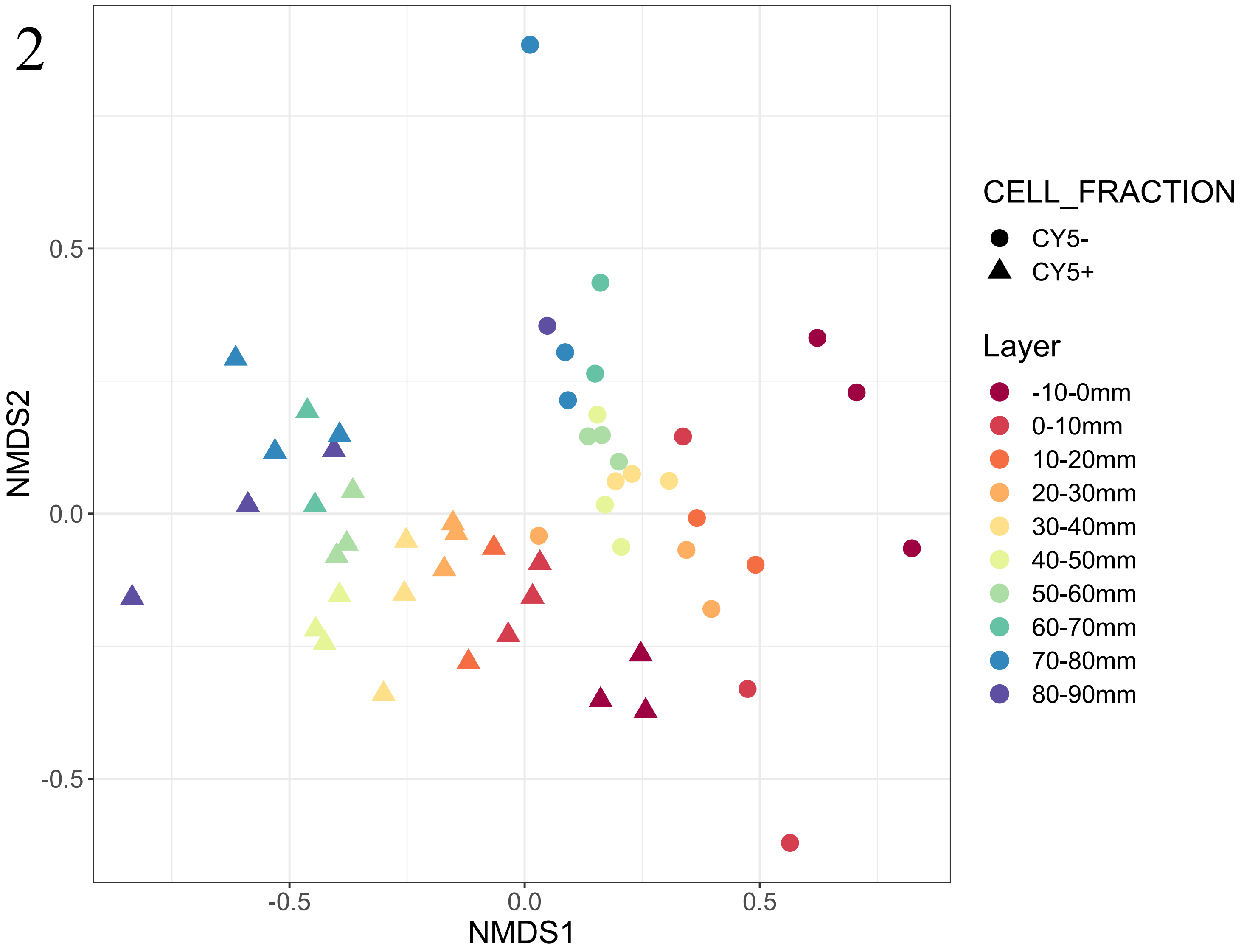

### Figure S5

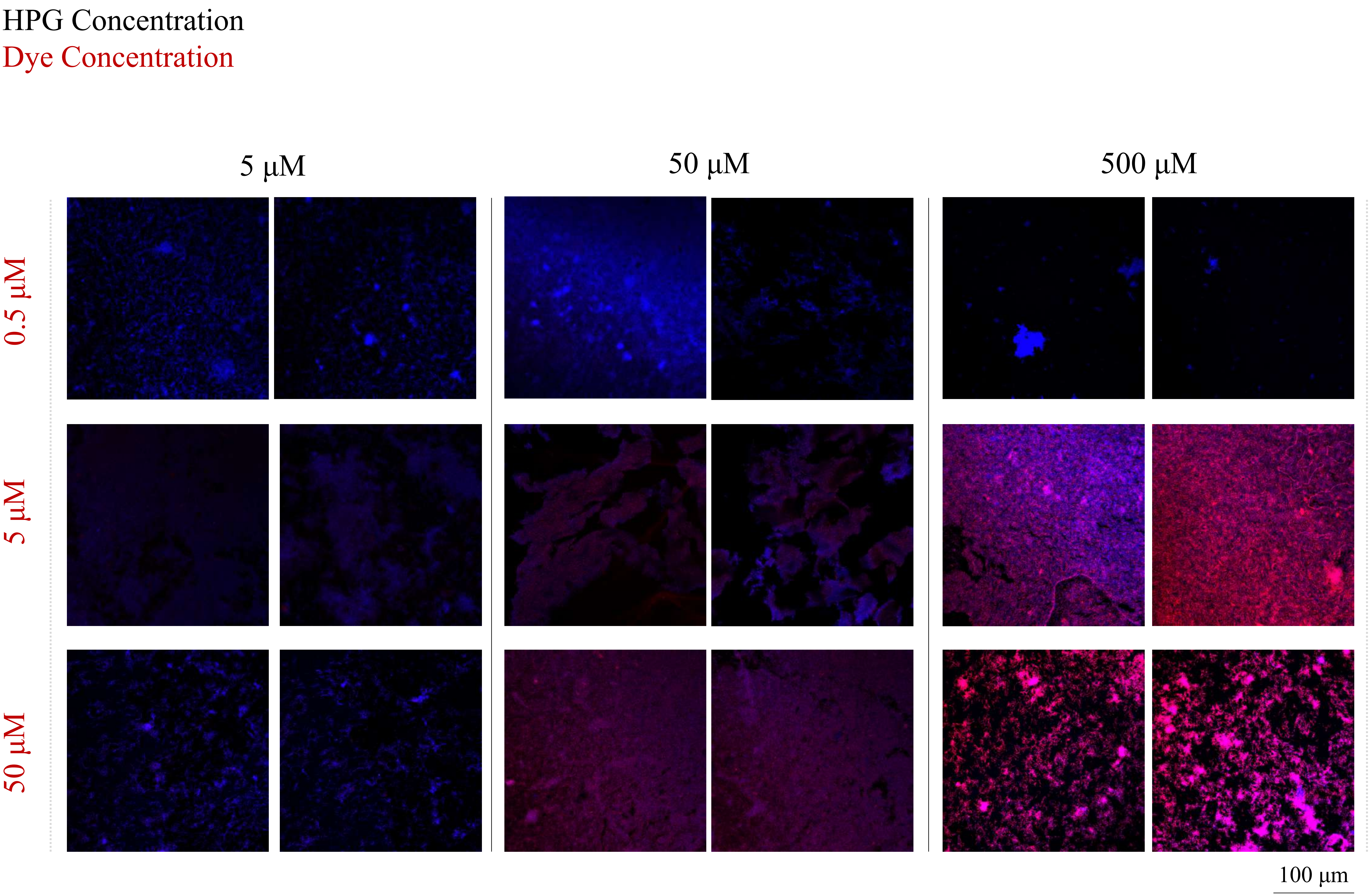

### Figure S6

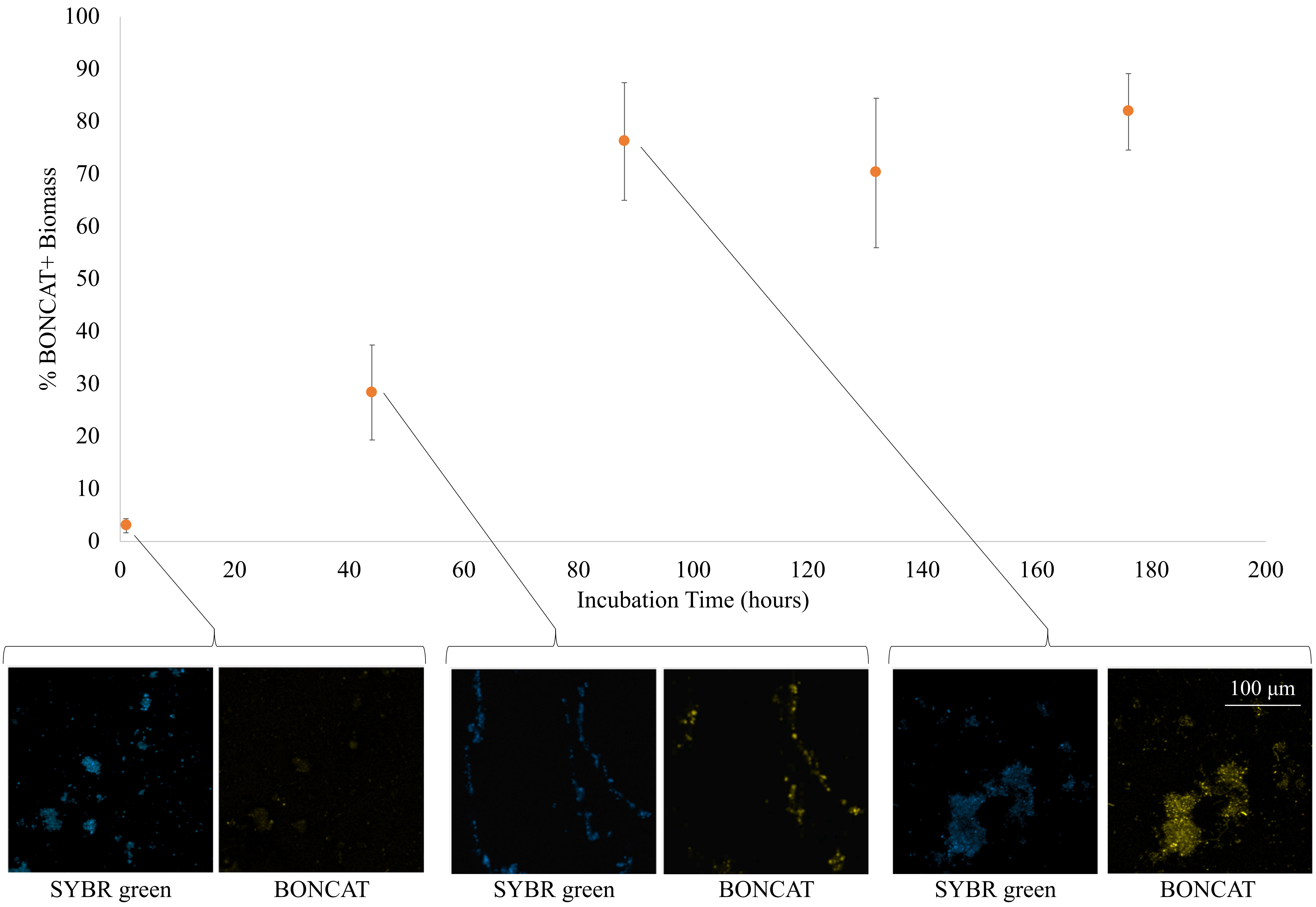

### Figure S7

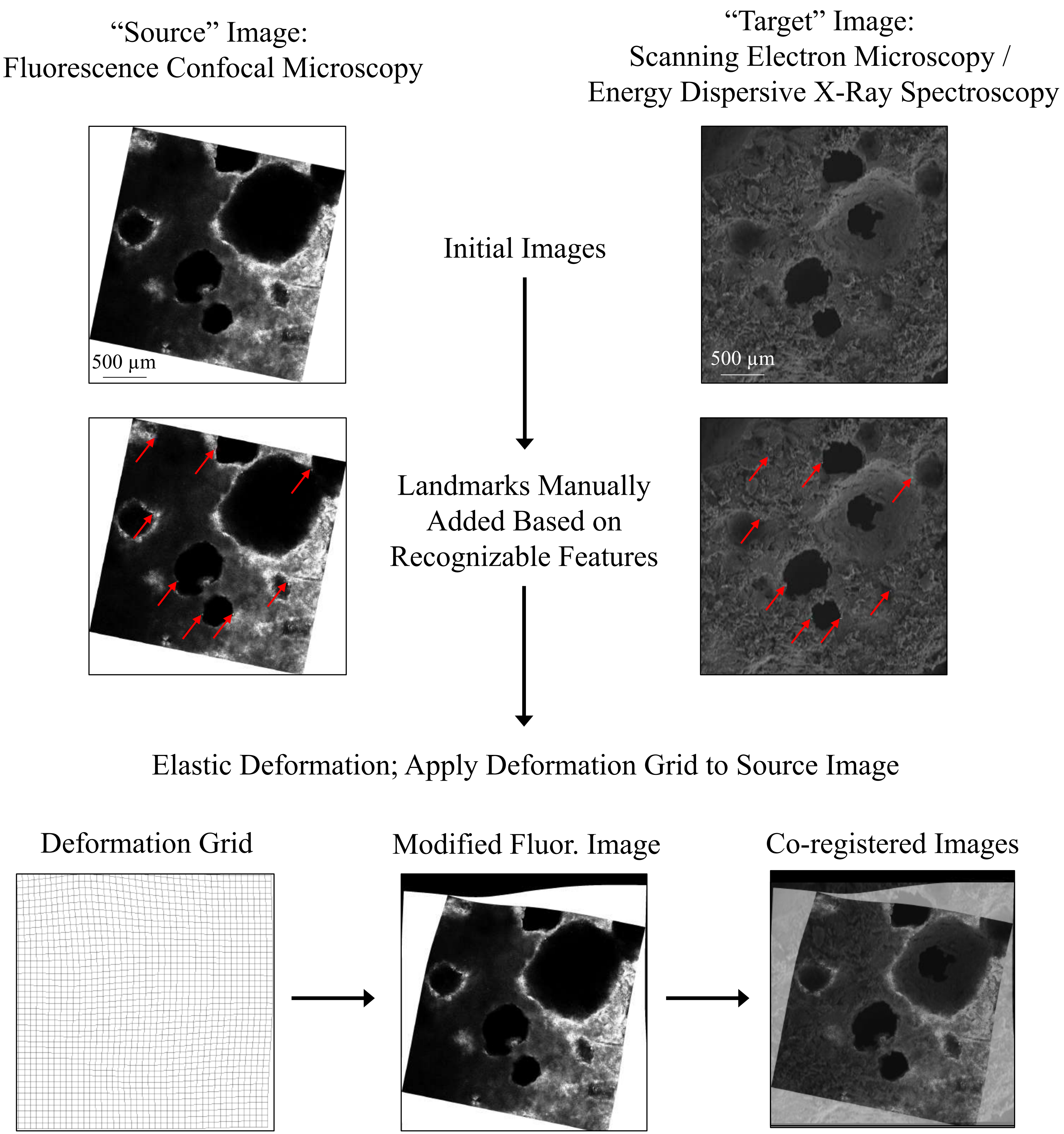

### Figure S8

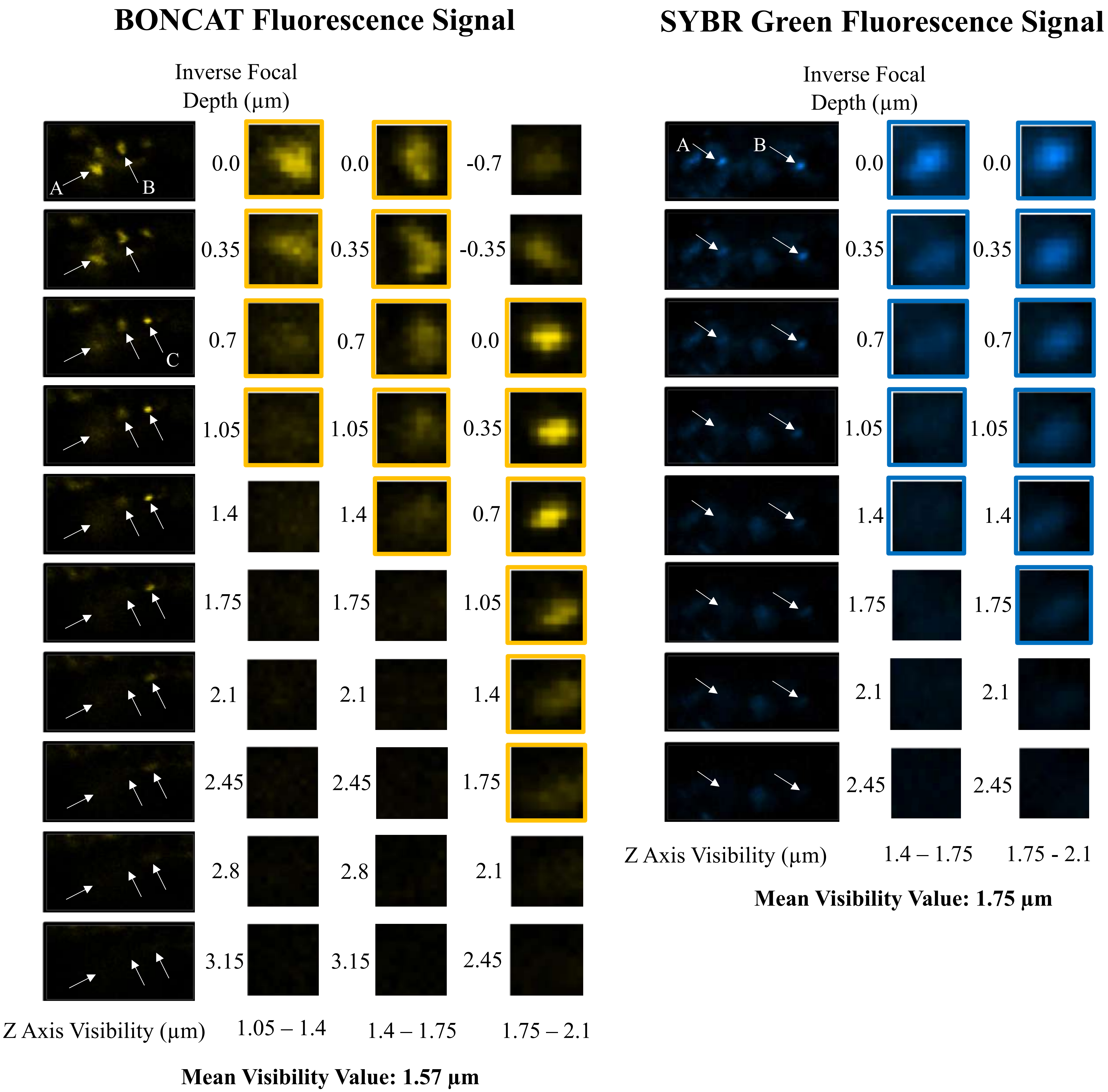

### Figure S9

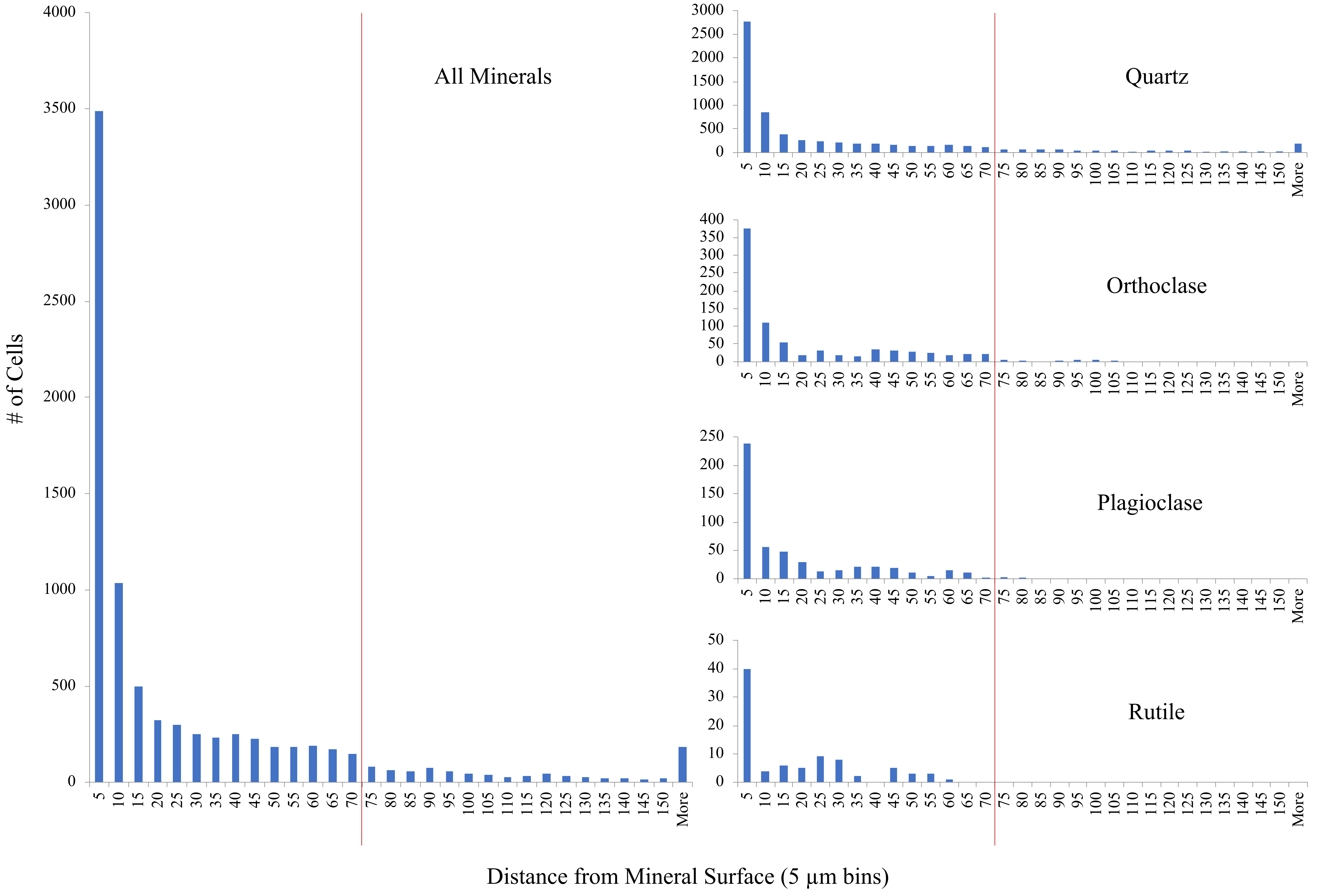

### Figure S10

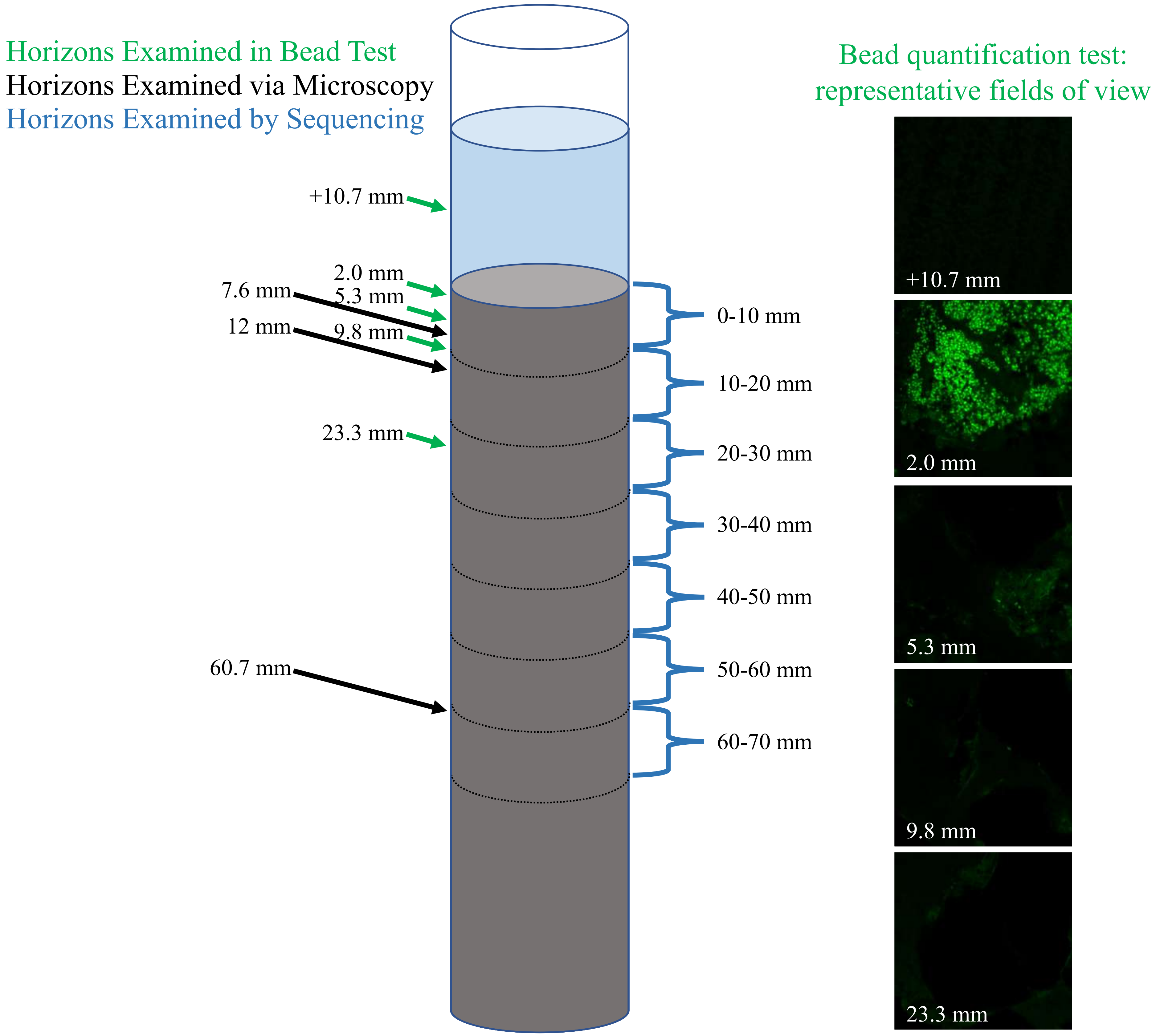
