## Supplementary material for "Spatially-resolved correlative microscopy and microbial identification reveals dynamic depth- and mineral-dependent anabolic activity in salt marsh sediment": Table S1

Table S1: Details on the conditions and analyses to which experimental and control sediment cores were subjected.

| **Sample Name** | **HPG** | **Autoclave-Sterilized** | **Fluorescence & Electron Microscopy** | **Cell Sorting & Sequencing** |
|---|---|---|---|---|
| BM | 50 µM | No | X | |
| BS | 50 µM | No | | X |
| CM | - | No | X | |
| CS | - | No | | X |
| AM | 50 µM | Yes | X | |

*Sample AM was homogenized sediment from 0-10 cm depth, incubated under lab conditions at room temperature.
